# Actuation Enhances Patterning in Human Neural Tube Organoids

**DOI:** 10.1101/2020.09.22.308411

**Authors:** Abdel Rahman Abdel Fattah, Brian Daza, Gregorius Rustandi, Miguel Angel Berrocal-Rubio, Benjamin Gorissen, Suresh Poovathingal, Kristofer Davie, Xuanye Cao, Derek Hadar Rosenzweig, Yunping Lei, Richard Finnell, Catherine Verfaillie, Maurilio Sampaolesi, Peter Dedecker, Hans Van Oosterwyck, Stein Aerts, Adrian Ranga

## Abstract

Tissues achieve their complex spatial organization through an interplay between gene regulatory networks, cell-cell communication, and physical interactions mediated by mechanical forces. Current strategies to generate in-vitro tissues have largely failed to implement such active, dynamically coordinated mechanical manipulations, relying instead on extracellular matrices which respond to, rather than impose mechanical forces. Here we develop devices that enable the actuation of organoids. We show that active mechanical forces increase growth and lead to enhanced patterning in an organoid model of the neural tube derived from single human pluripotent stem cells (hPSC). Using a combination of single-cell transcriptomics and immunohistochemistry, we demonstrate that organoid mechanoregulation due to actuation operates in a temporally restricted competence window, and that organoid response to stretch is mediated extracellularly by matrix stiffness and intracellularly by cytoskeleton contractility and planar cell polarity. Exerting active mechanical forces on organoids using the approaches developed here is widely applicable and should enable the generation of more reproducible, programmable organoid shape, identity and patterns, opening avenues for the use of these tools in regenerative medicine and disease modelling applications.

## Introduction

Generating complex and functional tissues during embryonic development relies on the action of mechanical forces that impose material deformations leading to precisely coordinated changes in tissue shape and identity^1-5^. These changes in shape and mechanical state serve critical and time-sensitive functional roles, for example in specifying cell fate^6,7^, spatially positioning stem cells^8^ and physically distorting the shape of morphogen fields^9^. Mechanical forces play a particularly significant role in neurulation, when an initially flat neural plate is shaped into a pseudostratified epithelial neural tube. The formation of the neural tube initiates by the furrowing of the neural plate, followed by bending at the single median hinge point and at the paired dorsolateral hinge points, and final apposition and fusion of the neural folds along the axis^10^. Neurulation requires both intrinsic forces generated within the neural plate by the apical constriction of cells at the midline, and extrinsic forces originating from tissues adjacent to the neural plate^10-12^. These extrinsic forces are driven by surface epithelium lying laterally to the neural folds and induce neural plate bending through a buckling mechanism. However, it is still unclear how neural tissues integrate mechanical cues to coordinate fate specification with patterning, growth and morphogenesis, and how mechanical cues and cell-intrinsic gene regulatory networks are linked^13^.

Organoids are ideal model systems to study the emergence of multicellular tissue complexity, as they recapitulate key elements of in-vivo development. Current methods for building organoids usually start with structures made by cellular aggregation^6,14^ or bioprinting^15-18^ to impose a static form, and then rely on self-organization to direct cells to their correct identity and place. To design supportive 3D microenvironments for organoid culture, an important focus in recent years has been on engineering the stiffness, degradability and extracellular matrix (ECM) composition of synthetic extracellular matrices^19,20^. The functionality of such matrices has been augmented by the addition of features such as hydrolytically or light-mediated degradation profiles^21-23^, and platforms have been developed to explore the role of each of these features using artificial 3D extracellular arrays^24,25^. Although such engineered matrices have helped in supporting organoid growth, they have proven limited in guiding patterning and morphogenesis as they are unable to provide active mechanical forces in a temporally defined manner, as occurs in-vivo. Indeed, attempts to generate shape-changing biological structures in-vitro have largely focused on actuating cell monolayers. For example, contractile canine kidney (MDCK) cells have been engineered to pull on a collagen substrate to generate origami-like folds^26^ in a cell type-specific method relying on cell-mediated tractions which cannot be temporally specified or controlled.. In the context of modeling neural development, microfluidic ligand gradients have been used to generate millimeter-scale patterns in hPSC-derived cell monolayers^27,28^, and neuroepithelial cells have been made to adhere to mechanically activated micropatterns leading to perturbed patterns of gene expression^29^. These techniques have remained confined to 2D however, with limited flexibility in actuation mode. The importance of mechanical perturbation on 3D tissue has also been shown in an *in vivo* context, where intestinal organoids were shown to mature when transplanted into the mesentry of mice and loaded via a spring-based system^30^. It has nonetheless not been possible to actuate organoids in-vitro, and studying how active mechanical forces regulate organoids in a controlled, developmentally biomimetic context has remained challenging. Here, we develop a platform that can provide active mechanical force to guide the fate, patterning and morphogenesis of organoids in 3D..

### Stretching enhances floor plate patterning

In order to understand the role of mechanical forces on early human neural fate specification and patterning, we reasoned that a patternable 3D organoid model of neurulation would be necessary to deconstruct and manipulate the relationship between cells and forces. Morphogens such as SHH, RA, BMP4 and WNT play a determining role in specifying cellular identities in the developing neural tube^31^. We have previously shown that RA alone is sufficient to pattern mouse neural tube organoids (mNTOs) derived from single mouse embryonic stem cells, but that competency to these signals is temporally restricted between the second and third day of differentiation^25^. To assess whether these conditions could be used to generate hPSC-derived neural tube organoids (hNTOs), single hPSC cells were embedded in a synthetic poly(ethylene) glycol (PEG)-based extracellular matrix whose composition and stiffness matched previously optimized properties for mNTOs. In combination with RA, we supplemented the growth medium with smoothened agonist (SAG) for floor plate (FP) induction, as previously shown^32^. These conditions were permissive for the growth of multicellular pseudostratified epithelial organoids (**Fig. 1a and Supplementary Fig. 1a**). We assessed neural patterning in these organoids at day 11 by immunohistochemistry for FOXA2, a marker of the ventral-most FP domain, and found that the optimal competence window for FP induction and patterning for hNTOs occurred later than in mNTOs, between day 3 and day 5, consistent with reports of delayed differentiation between mouse and human in-vitro development^33^ (**Fig. 1a and Supplementary Fig. 1b**). Patterning was assessed using a combination of FOXA2 area ratio (AR) and FOXA2 intensity (see methods).

**Fig. 1.**
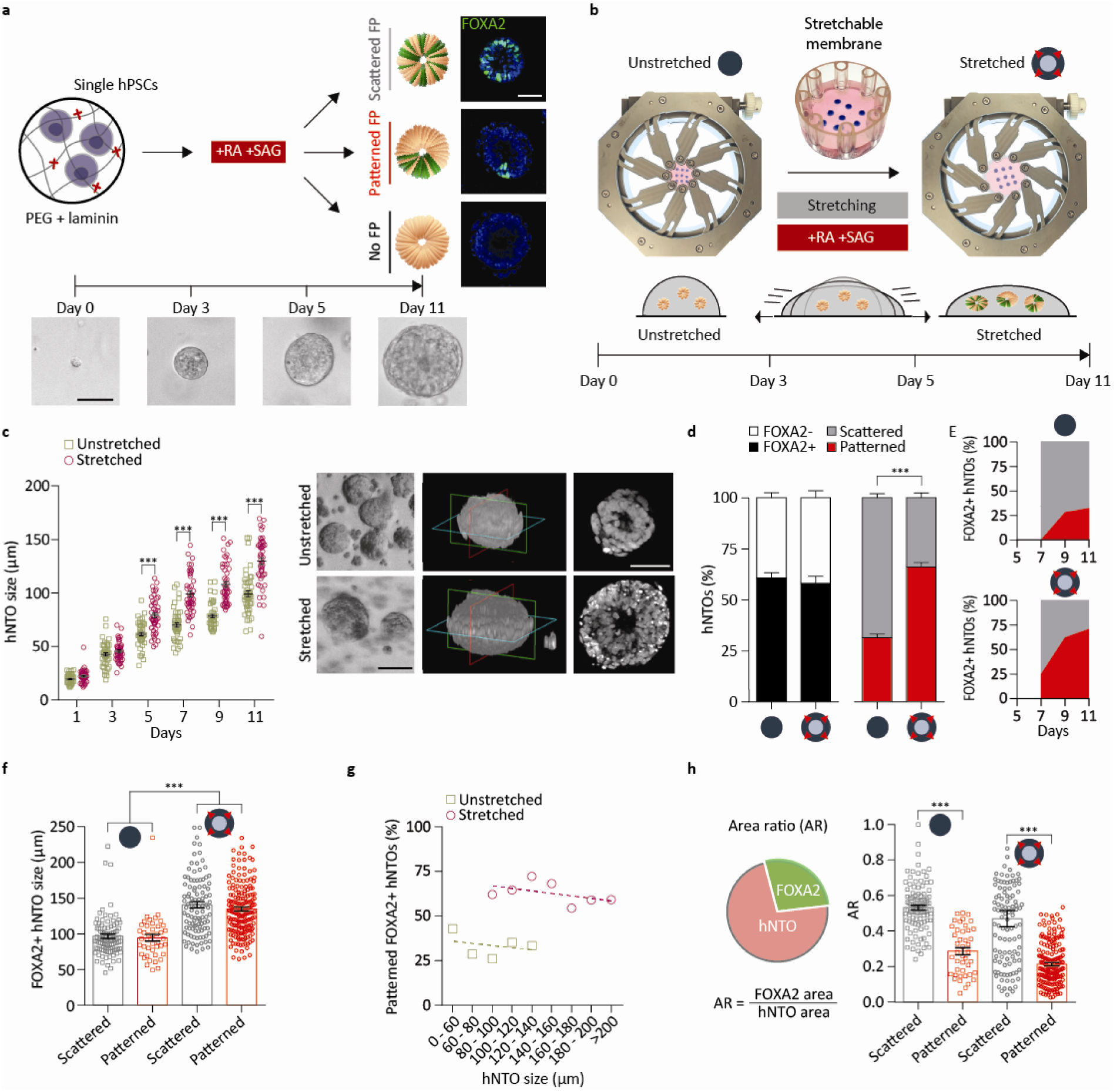
Actuation of hNTOs enhances organoid growth and patterning. **a** Schematic representation of the hNTO differentiation protocol. Single hPSCs are embedded into non-degradable PEG hydrogels supplemented with laminin. Treatment with RA and SAG induces the floor plate domain. Images show hNTO size at days 0, 3, 5 and 11, with three possible FP expression outcomes categorized based on scattered, patterned or absent FOXA2 expression. **b** Actuation of hNTOs. Hydrogel droplets (shown in blue for visualization) are placed on a stretchable membrane attached to the arms of an equibiaxial actuation device. Stretching occurs simultaneously with RA-SAG treatment (days 3-5), increasing membrane surface area by 100% at day 5 and maintained thereafter. Stretch-induced strains transferred to hydrogels modify the mechanical state of resident hNTOs. **c** Stretching increases hNTOs size starting at day 5 (50 hNTOs per data point), also seen in day 11 representative brightfield images, 3D confocal reconstructions and corresponding optical sections. **d** Quantification of FP induction and FP pattering events in FOXA2+ hNTOs (n = 5 for unstretched (303 hNTOs) and n = 6 for stretched (480 hNTOs) conditions). **e** Quantification of FOXA2 dynamics from day 7 (n = 2 for a total of >20 hNTOs per day). **f** Quantification of organoid size for stretched and unstretched conditions, binned for scattered and patterned hNTOs (hNTOs from d). **g** Patterned hNTO frequency as a function of binned organoid size for stretched and unstretched conditions (hNTOs from d), **h** Area ratio (AR) quantifying the ratio between FOXA2+ domain area and total organoid area, for scattered and patterned hNTOs within each condition (hNTOs from d). Error bars are SEM, *** p < 0.001, scalebars 50 μm.

To apply extrinsic mechanical forces on these organoids, we designed a process that enabled the programmable actuation of gel-embedded hNTOs. Taking advantage of the adhesion between PEG-based hydrogels and silanized silicone, we transferred strains from a stretchable silicone membrane to growing organoids within cross-linked PEG using an instrument equipped with programmable equibiaxially-actuated arms (**Fig. 1b**). As mechanical forces during the initiation of neural tube bending and closure occur simultaneously with patterning induction via notochord-derived signaling^34^, we reasoned that the initiation of organoid actuation could occur at the same time as morphogen treatment. The actuation of organoids was therefore initiated at day 3, gradually increased to 100% area (41% radial strain) by day 5 and was thereafter maintained until the experiment endpoint at day 11. Actuation resulted in a 30% increase in organoid size compared to unstretched controls (**Fig. 1c**). The number of cells per organoid was proportional to organoid size, demonstrating that the observed increase in size was due to higher proliferation rather than to a decrease in cell density (**Supplementary Fig. 2a-b**).

In order to understand the role of stretching on organoid differentiation and patterning, the number of organoids containing cells with FOXA2+ floor plate identity, as well as those exhibiting a patterned FOXA2+ domain was quantified. The proportion of organoids in which FOXA2+ cells could be detected was approximately 60% in both conditions, indicating that stretching did not alter the incidence of cells with floor plate identity. Strikingly, actuation enhanced the proportion of organoids with a patterned FOXA2+ domain, from 25% in the unstretched control (303 hNTOs) to 60% in the stretched condition (480 hNTOs) (**Fig. 1d and Supplementary Fig. 3**). In both conditions, FOXA2 expression was first observed 2 days after the end of the RA-SAG treatment. FOXA2 expression was initially scattered, but subsequently FP patterning increased, particularly for stretched hNTOs (**Fig. 1e**). These results suggest that mechanical forces play an important role in specifying spatial relationship between cells in early hNTO development.

In order to determine whether stretching biased hNTOs via a size-dependent mechanism, we quantified scattered versus patterned organoid sizes in each condition. Whereas organoids were on average larger in stretched conditions, there were no significant size differences within each condition between organoids with scattered or patterned FOXA2 expression (**Fig. 1f**). Moreover, the increased frequency of FP patterning upon stretch was maintained across hNTO sizes (**Fig. 1g**), indicating that, within the tested conditions, organoid size does not determine patterning frequency. The patterning of the domains within the neural tube has been reported to scale with growing tissue size^35^ and between species^36^. In order to explore whether these observations could be recapitulated with hNTOs we quantified the proportion of the projected organoid area occupied by the floor plate domain (**Fig. 1h**). For patterned organoids, the FOXA2+ domain represented approximately 20% of the total area, whereas in unpatterned organoids, FOXA2 expression covered 50% of the total. These observations were maintained across size and stretch conditions, suggesting that FP patterning in hNTOs is a scalable process as in-vivo, and that organoid stretching enhances patterning frequency without disrupting pattern scalability.

### Actuation time and magnitude, along with matrix stiffness, modulate floor plate patterning

To gain insights into the role of force magnitudes and duration on hNTO patterning, we modulated the extent and time of hNTO exposure to stretch. FP patterning efficiency increased with stretch magnitude and decreased when initiation of stretch was delayed, or by early removal from the stretching device (**Fig. 2a**). In particular, we observed a linear dependency on stretch magnitudes and a mechanical temporal competence window between days 3-7 when mechanical stimulation had the greatest and irreversible effect on enhancing FP patterning. These observations suggest that hNTOs retain a mechanical memory of applied forces, in line with emerging experimental^37^ and theoretical^38^ results in other systems. To test whether stretch direction would bias the direction or frequency of patterning, we built a device which allowed us to stretch organoids uniaxially at the same rate as that applied with the equibiaxial device. Increased FP patterning was observed in uniaxially stretched hNTOs, similarly to organoids stretched equibiaxially, however no preferential patterning orientation was observed, indicating that force magnitude rather than directionality is involved in the enhanced hNTO patterning phenomena. (**Supplementary Fig. 4a-f**).

**Fig. 2.**
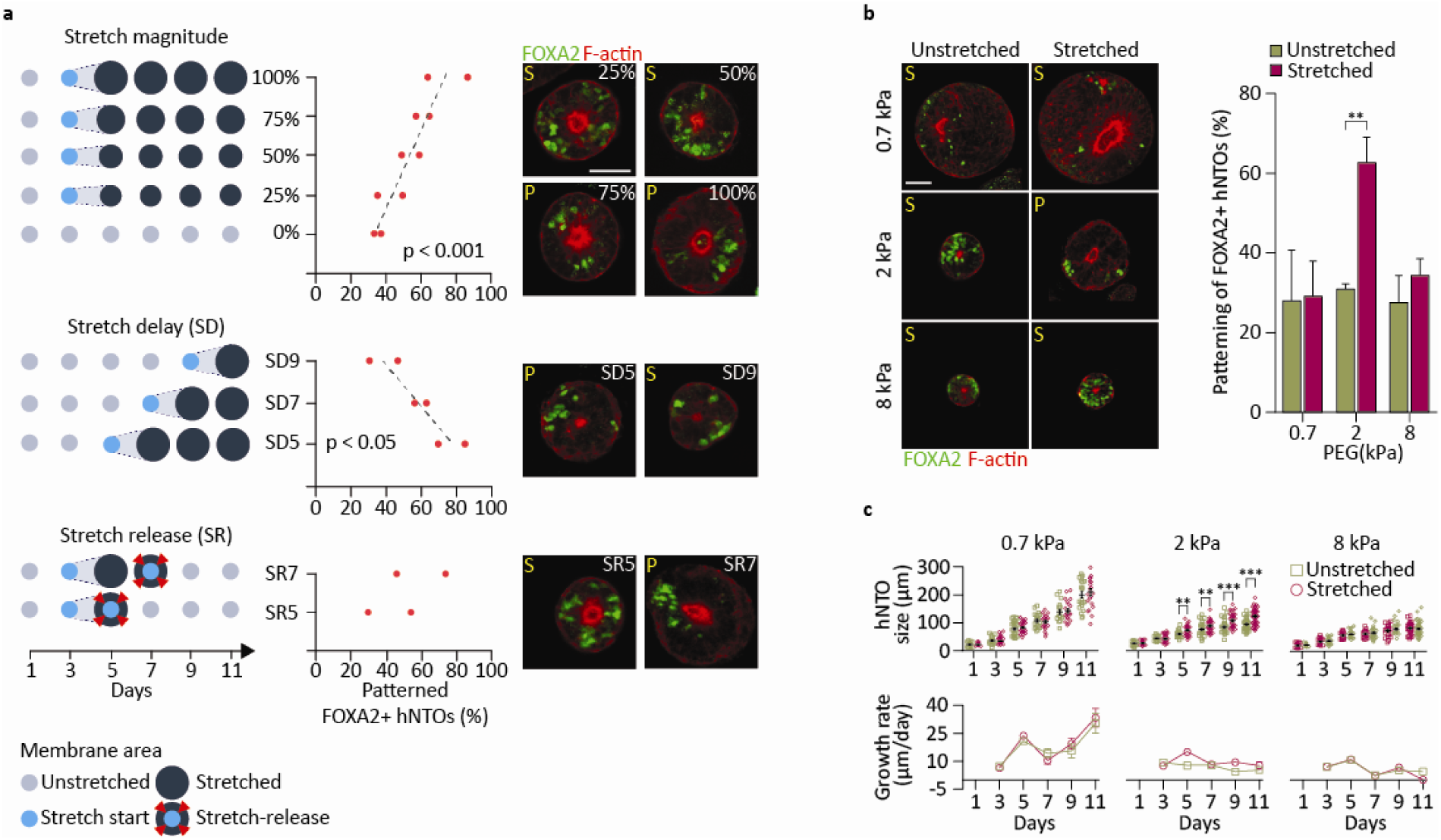
Floor plate patterning modulation by stretch magnitude, timing and interaction with matrix stiffness. **a** Diagrammatic representations of stretch magnitude regime modulation from 0% (no stretch) to 100% (maximal stretch, in area increase) as well as stretch delay and stretch release modulation experiments, with corresponding quantification of patterning efficiencies and representative images showing FOXA2 and F-actin expressions. (Error bars are SD, n = 2 for a total of > 60 hNTOs for all data points except for stretch magnitudes 75% (37 hNTOs) and 25% (34 hNTOs)). S and P denote scattered and patterned FP assignment respectively. P-values denote Pearson correlation analysis. **b** FOXA2 and F-actin expressions in hNTOs embedded in soft (0.7 kPa), intermediate (2 kPa) and stiff (8 kPa) matrices. Quantification of stretch-induced FP patterning for each matrix stiffness condition (n = 3 for unstretched 0.7 kPa (84 hNTOs), 2 kPa (47 hNTOs), 8 kPa (164 hNTOs) and stretched 0.7 kPa (58 hNTOs), 2 kPa (73 hNTOs), 8 kPa (152 hNTOs)). **c** Quantification of organoid sizes and growth rates in three matrix stiffnesses, with and without stretch (>30 hNTOs per data point except for 0.7 kPa (> 15 hNTOs)). Error bars are SEM unless stated otherwise, ** p < 0.01, *** p < 0.001, scalebars 50 μm.

We and others have previously shown that the stiffness of the local microenvironment can modulate organoid patterning and morphogenesis^23,25^. To deconvolve the role of active (actuated) versus passive (matrix) mechanics on FP patterning, we stretched hNTOs embedded in soft (0.7 kPa), intermediate (2 kPa) and stiff (8 kPa) PEG hydrogel matrices. FP patterning efficiency was only enhanced at intermediate matrix stiffness and remained unchanged upon stretching in soft and stiff matrices (**Fig. 2b**). In parallel, we observed that while organoids increase in size at the fastest rate in the softest matrix, a relative increase in size upon stretch was only evidenced at intermediate 2 kPa stiffness (**Fig. 2c**). These results suggest a dynamic mechanical feedback whereby organoid growth compresses the adjacent matrix, with the concomitant matrix resistance counteracting organoid growth and that the same feedback is likely involved in pattering enhancement upon stretch.

To interrogate the effects of such changes to the mechanical microenvironment on organoid growth in actuated conditions, we assessed matrix strain in the direction of stretch (ε_rr_). A simplified stress measure (σ_rr_) was then calculated by multiplying the measured strain by the matrix elastic modulus (**Supplementary Fig. 5a, Fig. 3a**, see methods). In unstretched conditions, we observed matrix compression (ε_rr_ < 0, σ_rr_ < 0) driven by hNTO growth. In matrix regions devoid of hNTOs, stretching generated tensile strains and stresses (ε_rr_ > 0, σ_rr_ > 0). Finite element analysis simulations of equibiaxial stretching further confirmed that the generated tension field was homogeneous throughout most of the matrix volume, with the exception of the outermost edge regions of the hydrogel, which in experiments is largely devoid of hNTOs (**Supplementary Fig. 5b**). In matrix regions containing hNTOs, organoid growth-induced compression was greater than stretch-induced tension, leading to the conclusion that actuation did not impose net tension stresses on hNTOs (**Fig. 3a**). Instead, stretching provided relief of growth-induced compression in the vicinity of the hNTOs, such that ε_rr,stretched_ > ε_rr,unstretched_ and σ_rr,stretched_ > σ_rr,unstretched_ for days ≥ 5 (**Fig. 3a**). To investigate whether the magnitude of compressive stresses drive organoid growth, we analyzed the relationship between σ_rr,Day11_ and hNTO growth rates. A fast (> 10 μm/day) and a slow (< 10 μm/day) growth regime were observed, which were separated by a critical stress σ_rr,critical_ = −0.81 kPa (**Fig. 3b**).

**Fig. 3.**
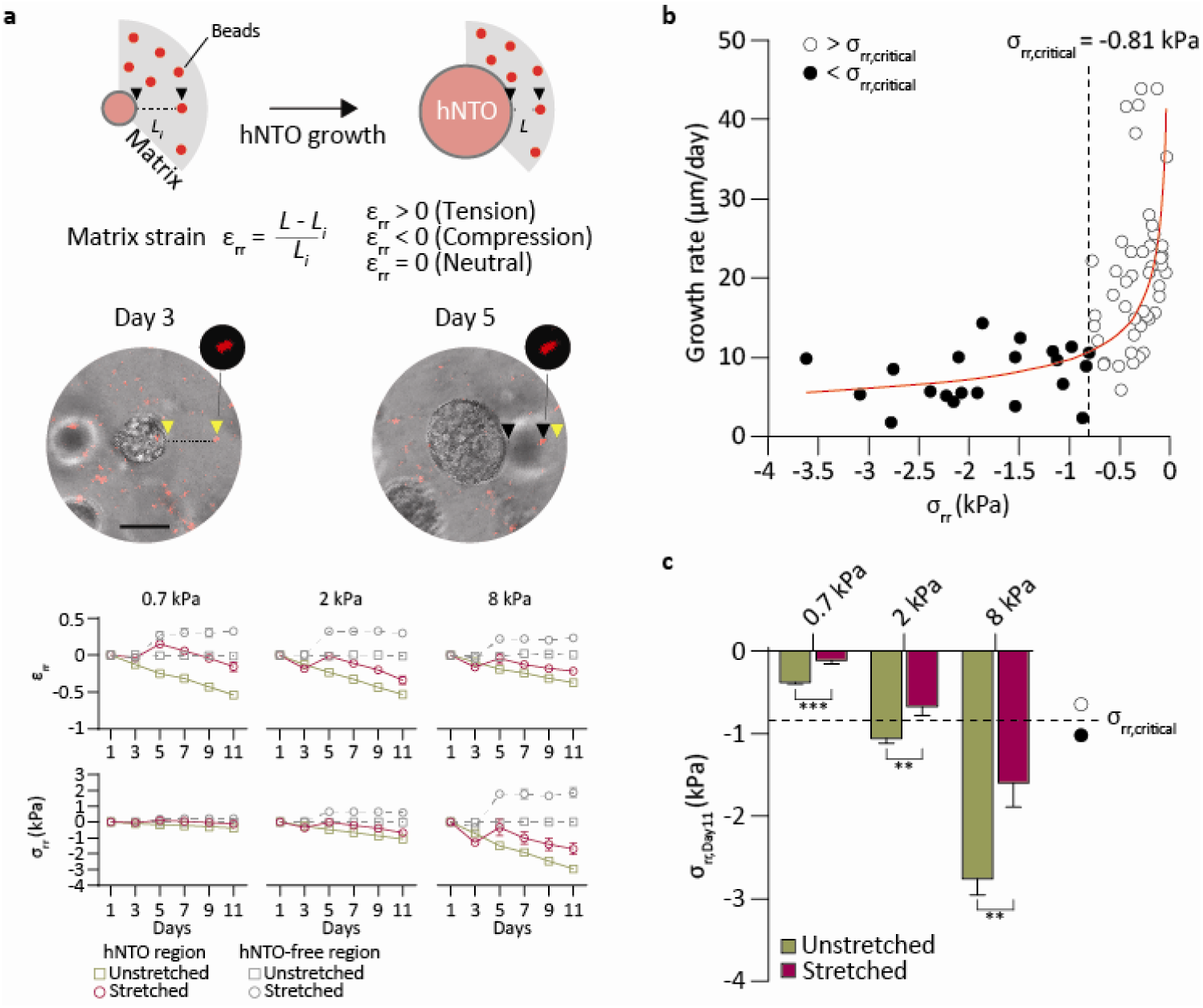
Matrix actuation modulates mechanical microenvironment by relieving growth-induced compression. **a** Schematic representation of bead tracking technique, corresponding representative images and quantification of matrix radial strain and stress for soft, intermediate and stiff matrix conditions (15 segments per datapoint). **b** hNTO growth rates and corresponding matrix compressive stress states define two main growth behaviors 1) fast growth rate for σ_rr_ > σ_rr,critical_ and 2) slow growth rate for σ_rr_ < σ_rr,critical_ where σ_rr,critical_ = −0.81 kPa. Red curve represents best fit power curve (y = 193.66(-x)^-0.432^, R^2^ = 0.4788). **c** Compressive stress states at different stiffnesses (from **Fig. 2c**). Soft and hard matrices remain in fast (σ_rr,Day11_ >> σ_rr,critical_) and slow (σ_rr,Day11_ << σ_rr,critical_) growth regimes respectively regardless of stretch condition, with only intermediate stiffness allowing for a growth regime transition from slow (σ_rr,Day11_ < σ_rr,critical_) to fast (σ_rr,Day11_ > σ_rr,critical_) upon stretch. Error bars are SEM, ** p < 0.01, *** p < 0.001, scalebars 50 μm.

Because soft matrices exert little counteracting pressure on organoid growth, σ_rr,Day11_ >> σ_rr,critical_ in both stretched and unstretched conditions (**Fig. 3c**) and hNTOs therefore have similarly high growth rates and large sizes (**Fig. 2c**). Conversely, in stiff matrices, although actuation relieves significant stress, both conditions result in σ_rr,Day11_ << σ_rr,critical_, and thus hNTOs have similar low growth rates resulting in small hNTOs. In contrast to soft and stiff matrices, unstretched intermediate matrices have σ_rr,Day11_ < σ_rr,critical_, corresponding to slow growth conditions. Upon stretch, a transition to the faster growth regime occurs where σ_rr,Day11_ > σ_rr,critical_, resulting in faster growth rates (**Fig. 2c**). High growth rates do not by themselves impose higher patterning frequency since low frequencies of patterning are observed in organoids with highest growth rates in soft matrices. However, a transition from low to high growth rate regimes, only possible in matrices of intermediate stiffness, appears necessary for enhancing FP patterning. This suggest that a matrix stiffness matching that of the in-vivo microenvironment is not only permissive to growth and cellular reorganization, but also functions as a substrate for optimal transfer, at high fidelity, of extrinsic forces that lead to FP patterning enhancement. Moreover, the growth rate transition in actuated intermediate stiffness matrices likely alters specific cellular programs responsible for increasing FP patterning frequencies.

### Transcriptional landscape of actuated hNTOs

To comprehensively define the cellular identities in hNTOs and to unravel underlying differences between control and stretched hNTOs, we performed scRNAseq on organoids at three time points: before stretch (day 3), immediately after reaching final stretch (day 5) and after maturation (day 11). Graph based clustering of the 17,826 cells retained for analysis indicated increased cluster diversity over differentiation time (**Fig. 4a**) which was confirmed by pseudotime analysis (**Supplementary Fig. 6**). We identified 11 clusters in this dataset, and annotated them to reveal groups of cells with dorsal (D-11), intermediate (I-11), ventral (V-5, V-11), forebrain (FB), neuroprogenitor (NP-5, NP-3), neural crest (NC), neural crest derivatives (NCD) identities, as well as two groups with shared identities assigned as transition clusters (T-a and T-b) (**Fig. 4a-c, Supplementary Fig. 7a-b, Supplementary table 1**). The transcriptomic signature of human-specific neural differentiation was confirmed by multiple markers (**Supplementary Fig. 8**), including the presence of the earliest human neural marker *PAX6*^*39*^ observed in clusters corresponding to day 3 and day 5, and a transition to the more differentiated neuroepithelial marker *SOX1* seen by day 11 (**Fig. 4d**).

**Fig. 4.**
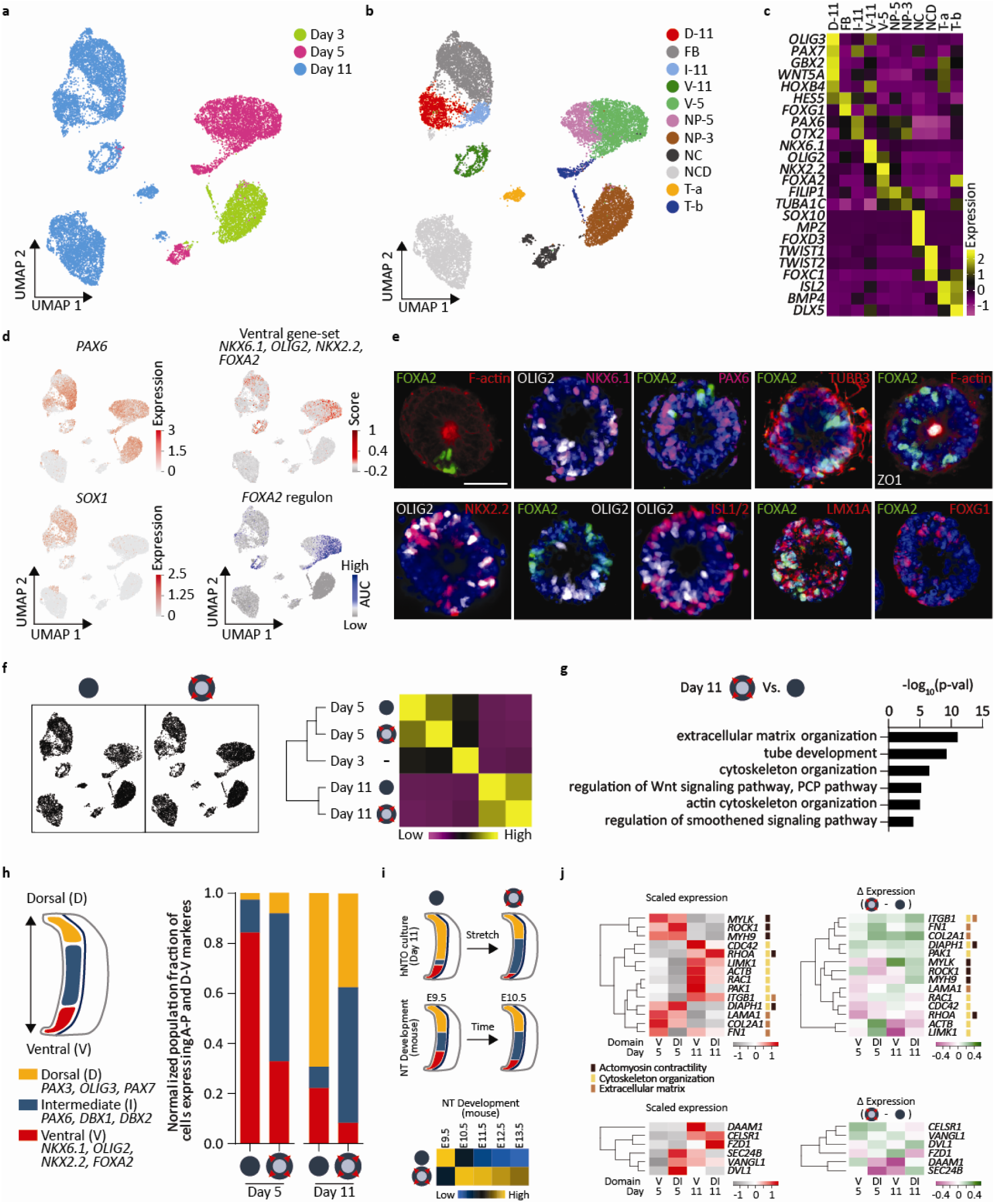
Single cell RNA-seq of hNTOs reveals actuation-specific transcriptomic changes. **a** UMAP of the combined dataset (day 3, with day 5 and day 11 stretched and unstretched samples). **b** UMAP of the combined dataset with identified clusters (Dorsal (D-11), forebrain (FB), intermediate (I-11), ventral (V-11), neuroprogenitors (NP-5 and NP-3), neural crest (NC), neural crest derivatives (NCD), and transition clusters (T-a and T-b). Top marker genes for each cluster and unannotated clusters in (**Supplementary table 1 and Supplementary Fig. 7a-b**). **c** Selected marker genes for anteroposterior, dorsoventral, neural crest, neural crest derivative identities are exclusively present in specific clusters. Expression values are normalized and centered. **d** UMAPs showing expression of *PAX6*, early neural and intermediate dorsoventral marker, present in multiple clusters, while *SOX1*, late neural progenitor marker, is expressed only in day 11 clusters. UMAP showing highest ventral gene-set scores in ventral clusters V-5 and V-11. AUC values confirm *FOXA2* regulon activity is highest in ventral and T-b clusters. **e** fate specification by IHC in unstretched hNTOs. (IHC for stretched hNTOs in **Supplementary Fig. 10**). **f** UMAPs for unstretched and stretched hNTOs, with correlation heatmap and hierarchical clustering. **g** GO enrichment analysis highlighting upregulation of key processes including ECM, actin and cytoskeleton organization and PCP upon stretch in day 11 dataset. **h** Dorsoventral cell fractions based on binned cell identities showing lower ventral fractions in stretched hNTOs compared to unstretched counterparts. **i** Comparison to mouse neural tube development. Diagrams display approximate dorsal, intermediate and ventral population fractions in stretched and unstretched hNTO cultures at day 11 and in mouse NT E9.5 and E10.5 as previously reported^42^. For mouse NT development the dorsal, intermediate and ventral progenitor domains are color coded respectively as yellow (roof plate to dp3), blue (dp4 to p2), and red (p1 to FP). The reduced ventral and dorsal population fractions at the expense of intermediate cells upon stretch are comparable to the transition from E9.5 to E10.5. Correlation matrix mapping hNTO dorsoventral identity to mouse neural tube development. **j** Gene expression comparison for unstretched samples for contractility, cytoskeleton organization, ECM and PCP markers, and their modulation upon stretch. Expression values are normalized and centered. Scalebars 50 μm.

To track the differentiation of cells in the ventral domains, we established a gene set including *FOXA2, NKX2*.*2, OLIG2 and NKX6*.*1*, which was present in ventral clusters at days 5 and 11 (**Fig. 4d)**. Single-cell regulatory network inference and clustering (SCENIC) analysis was performed on the dataset^40^, identifying 404 regulons representing TFs and their predicted transcriptional targets containing statistically significant TF binding sites in their cis-regulatory control elements (**Supplementary table 2**). *FOXA2* regulon activity was confirmed within ventrally associated clusters (**Fig. 4d, Supplementary Fig. 9a-b**).

In order to validate the identities determined by scRNAseq analysis and to spatially relate them to the ventral domain, we used endpoint immunohistochemistry (IHC) (**Fig. 4e)**. hNTOs displayed pV3 NKX2.2+, progenitor motor neuron (pMN) OLIG2+, and ventral NKX6.1+ cells in close proximity to the FP^41^. Basal regions expressed ISL1/2+ MN cells and the early neuronal marker TUBB3. PAX6, also associated with intermediate domains, was observed mainly in FOXA2-organoids. Midbrain ventral marker LMX1A was co-expressed with FOXA2, while forebrain marker FOXG1 was present in FOXA2-hNTOs, in accordance with the expected lack of floor plate in the forebrain. Additionally, ZO1 expression confirmed the presence of apical tight junctions and co-localized with actin cables characteristic of apico-basally polarized epithelial organoids.

Overall, the transcriptional signature of stretched and control samples was similar, both at early (day 5) and late (day 11) time points, (**Fig. 4f**), suggesting that the imposed mechanical stimulation does not lead to changes in differentiation or to the emergence of additional cell fate types. In line with scRNAseq analysis, analysis, all markers visualized by IHC that were present in control conditions (**Fig. 4e**) were also observed in stretched hNTOs (**Supplementary Fig. 10**). To determine the transcriptional changes occurring upon stretch, differential gene expression analysis was performed followed by Gene Ontology (GO) enrichment analysis between conditions at day 11 (**Fig 4g**). We identified extracellular matrix production, actin and cytoskeleton organization, as well as planar cell polarity (PCP) as processes upregulated in stretched samples, highlighting their role in stretch-mediated mechanotransduction and patterning regulation.

To explore transcriptional differences in a domain-specific manner, cells were binned according to their dorso-ventral (DV) gene-set scores and their proportions were evaluated at day 5 and 11 (**Fig. 4h**). The ventral fraction in stretched organoids was reduced by 2.6 and 2.7 fold at day 5 and 11 respectively, suggesting that patterning was accompanied by a decrease in the number of floor plate domain cells, in line with morphological observations (**Fig. 4h**). To evaluate whether these results matched with in-vivo development, we compared proportions of hNTO cells found in each DV domain with a published mouse neural tube scRNAseq dataset^42^. In day 11 hNTOs, stretch resulted in a reduction of the ventral domain, expansion of intermediate domain, and reduction of dorsal domain, similarly to trends observed during mouse neural tube development from E9.5 to E10.5 (**Fig. 4i**). Correlation analysis of hNTO fate proportions to that of the developing mouse neural tube revealed that unstretched hNTOs resembled the early E9.5 development stage, while stretched conditions were closer to the more developed E10.5/E11.5 neural tube. These results suggest that actuation-mediated acceleration of patterning may stabilize more physiological domain proportions in hNTOs.

Because actuation changed domain-specific cell proportions, we next explored whether stretch-induced upregulation of processes related to ECM production, cytoskeleton remodeling, contractility and PCP exhibited domain-specific differential regulation (**Fig. 4j**). Contractility genes such as *ROCK1, MYLK, MYH9, DIAPH1*, as well as ECM genes *LAMA1, COL2A1, FN1* were upregulated before FP emergence, suggesting that these have a role in pattern induction. Conversely, *RAC1, CDC42, PAK1, DIAPH1*, involved in cytoskeleton organization were most expressed in the ventral domain at day 11 and were also the genes for which expression was most upregulated in this domain upon stretching, suggesting a role in pattern stabilization. Significant changes in PCP gene expression were also observed as a function of time, domain and mechanical state. Downstream PCP effectors *DVL1* and *SEC24B* were most highly expressed immediately after stretch in the dorsal/intermediate domain whereas receptors *VANGL1, CELSR1* and *FZD1* were upregulated at day 11. Interestingly, *DAAM1*, which links PCP with the contractility machinery^43^ had highest expression in the ventral domain at D11, but its expression also decreased the most in this domain upon stretch. Overall, these transcriptomic results suggest that pathways involved in contractility and PCP are likely mediators of patterning enhancement in actuated hNTOs.

### Role of contractility and PCP in actuated hNTOs

In order to further investigate the role of contractility in floor plate induction and patterning we used the ROCK inhibitor Y-27632 (ROCKi) to perturb hNTO contractility and phosphorylated myosin light chain activity. Prolonged exposure to ROCKi reduced FP induction from 59% to 30% while patterning frequency of FOXA2+ hNTOs treated with ROCKi was insensitive to stretch (**Fig. 5a**). Lumen formation is a hallmark of apicobasal polarization in neuroepithelial cysts and is necessary for floor plate patterning in mNTOs^25^. We observed that the frequent perturbation of organoid lumens was a clear phenotypic effect of ROCKi on hNTOs. In contrast to the round lumen present in more than 90% of untreated organoids, only 55% of ROCKi-treated organoids presented a lumen, with most of these having an elongated shape (**Fig. 5b**) likely due to increased cytoskeleton fluidity resulting from impaired contractility. The importance of lumen formation was underscored by the observation that less than 5% of organoids without a lumen were able to form a floor plate (**Fig. 5c**). Reduced FOXA2 induction as a result of ROCKi treatment can therefore be attributed to reduced lumenogenesis. The lack of patterning enhancement in stretched ROCKi-treated hNTOs was not linked to defects in lumen formation, as these were not significantly different between ROCKi-treated stretched and unstretched samples (**Fig. 5b**).

**Fig. 5.**
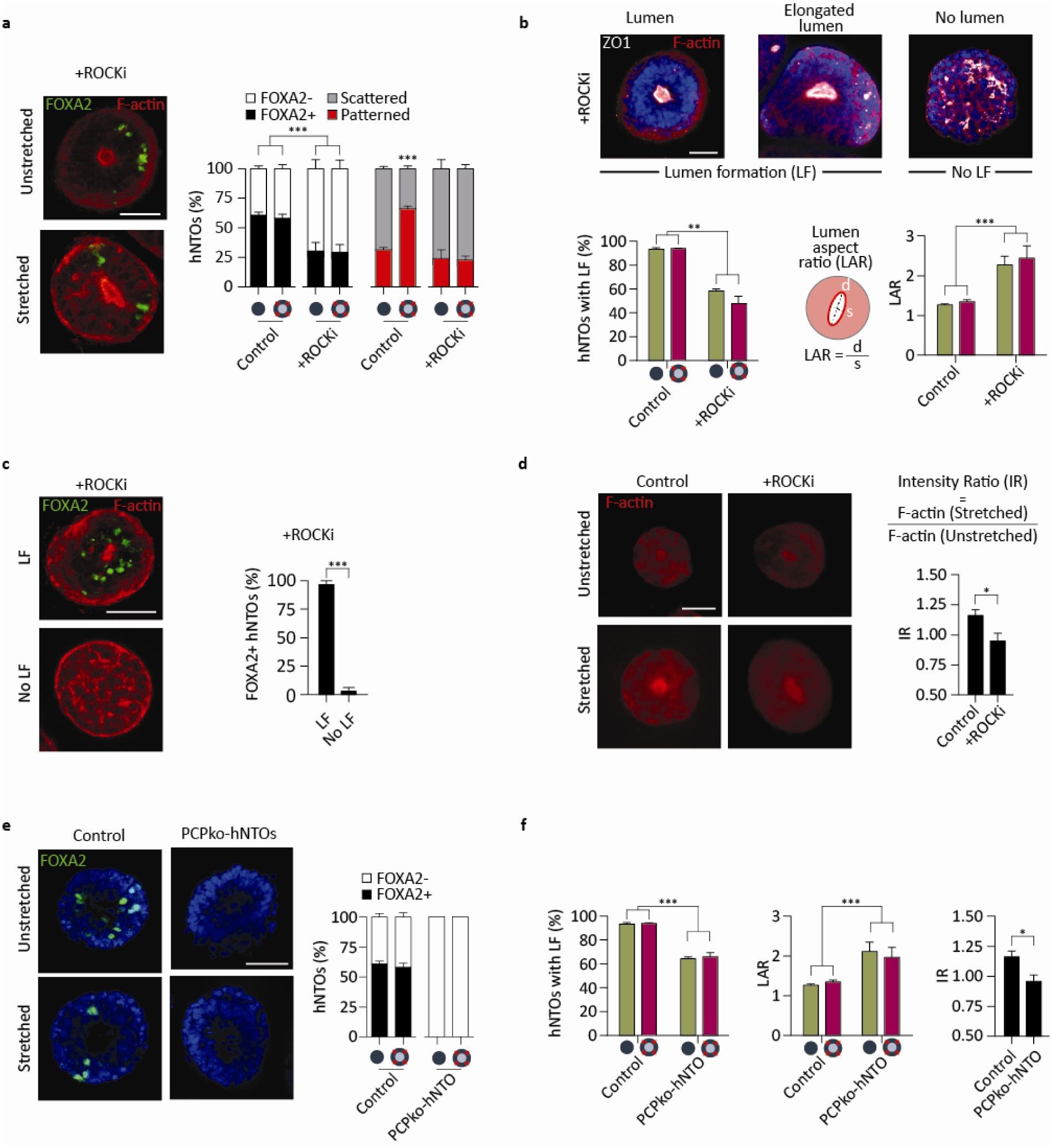
Contractility and planar cell polarity determine floor plate induction and hNTO patterning. **a** Representative images of FOXA2 and F-actin expression in stretched and unstretched hNTOs with prolonged ROCKi exposure, and quantification of FP induction patterning upon stretching compared to control conditions (n = 3 for unstretched (222) and stretched (187) hNTOs, control hNTOs from **Fig. 1d**). **b** Representative images showing F-actin and ZO1 expressions for hNTOs exposed to 10 μM of ROCKi for 11 days. Quantification of the number of hNTOs exhibiting lumen formation (LF) and lumen aspect ratio (LAR) (n = 3 for >25 hNTO per condition). **c** Representative images and quantification showing FOXA2 expression in hNTOs with and without LF (n = 3 for >30 hNTOs). **d** Representative hNTOs stained with F-actin, with quantification of F-actin intensity within the organoid. Intensity ratio (IR) quantifies the relative F-actin intensity within organoids between stretched and unstretched samples (n = 3 for >25 hNTO per condition). **e** Representative images highlighting the absence of FOXA2 expression in PCP^KO^-hNTOs compared to control hNTOs under unstretched and stretched conditions (n = 3 for >50 PCP^KO^-hNTOs, control hNTOs from **Fig. 1d**). **f** Quantification of patterning frequency, LAR and IR for PCP^KO^-hNTOs (n = 3 for >30 PCP^KO^-hNTOs per condition). Error bars are SEM, * p<0.05, ** p<0.01, *** p<0.001, scalebars 50 μm.

Cytoskeletal remodeling has been shown to increase under mechanical stimulation^44^, and we therefore hypothesized that it may be impaired upon ROCKi treatment. To test this, we evaluated the ratio of F-actin intensities within the hNTOs (intensity ratio, IR) between stretched and unstretched samples as a surrogate for remodeling activity. Stretched hNTOs had increased IR values (IR>1) compared to unstretched controls (**Fig. 5d**), while this phenomenon was completely abrogated upon ROCKi exposure (IR 1), outlining a mechanism of stretch-induced patterning enhancement involving contractility-mediated cytoskeleton remodeling.

The physical and molecular events occurring during neural tube morphogenesis must be coordinated with the concurrent convergent extension of the embryo. Planar cell polarity (PCP) signaling directly links these two processes, and had been identified as a differentially expressed pathway in our GO enrichment analysis. To investigate the links between PCP and floor plate patterning, we used a hPSC line with point mutations in PCP genes *CELSR1, VANGL1*, and *SEC24B*^45^, which have also been linked to neural tube defects^46-48^. The use of a triple mutant line ensures that PCP is completely abrogated. Unlike control and ROCKi-treated hNTOs, PCP^KO^-hNTOs exhibited the complete absence of FOXA2 (**Fig. 5e**) and high expression of PAX6 in both stretched and unstretched conditions (**Supplementary Fig. 11**). The complete lack of FP in PCP^KO^-hNTOs indicates that PCP plays an early role in FP induction, rendering the cells unresponsive to the RA-SAG patterning signal. Similar to ROCKi-treated hNTOs, PCP^KO^-hNTOs exhibited distorted cytoskeletons, marked by lower LFs, high LARs and unchanged IR values upon stretch (**Fig. 5f and Supplementary Fig. 12**), suggesting that PCP impairment reduced organoid-wide response to exogenous mechanical modulations. The cytoskeleton disorganizations observed in both ROCKi-treated and PCP^KO^ hNTOs and consequent lack of patterning enhancement in the former upon stretch suggest that the formation of organized epithelia is a critical requirement in hNTOs for responsiveness to patterning signals and their modulation by mechanical forces.

Altogether, our results demonstrate that organoids are responsive to exogenous mechanical modulation driven by matrix actuation. We show that human neural tube organoids have increased patterning events upon stretch through changes in their actomyosin contractility, cytoskeleton remodeling and PCP machineries (**Fig. 6**). We further show that this response depends on stretch magnitude and timing, as well as on an interplay with the extracellular matrix, whose stiffness must be tuned to optimally relay extrinsic forces. We identify parallels between the molecular behavior of actuated hNTOs and the morphological dynamics of in vivo neural tube development^12^. Our results suggest that mechanical forces in early human neural tube bending regulate the emergence of the floor plate and help orchestrate domain specification. The precise mechanisms leading to enhanced patterning upon actuation remain to be elucidated. It is possible that actuation confers developmental robustness and that fluctuations in contractility or PCP gene expression may be buffered by tissue-scale mechanical coupling, as has recently been shown during drosophila gastrulation^49,50^. The engineered platform we describe here to mechanically actuate 3D organoids, and the accompanying single-cell atlas, can be used in conjunction with synthetic matrices to completely specify the extrinsic mechanical state of organoids. Enlarging the toolbox of organoid biology to include actuation will allow for more robust and reliable 3D phenotypes, and will allow to interrogate the role of mechanotransduction on other developmental and disease model systems.

**Fig. 6.**
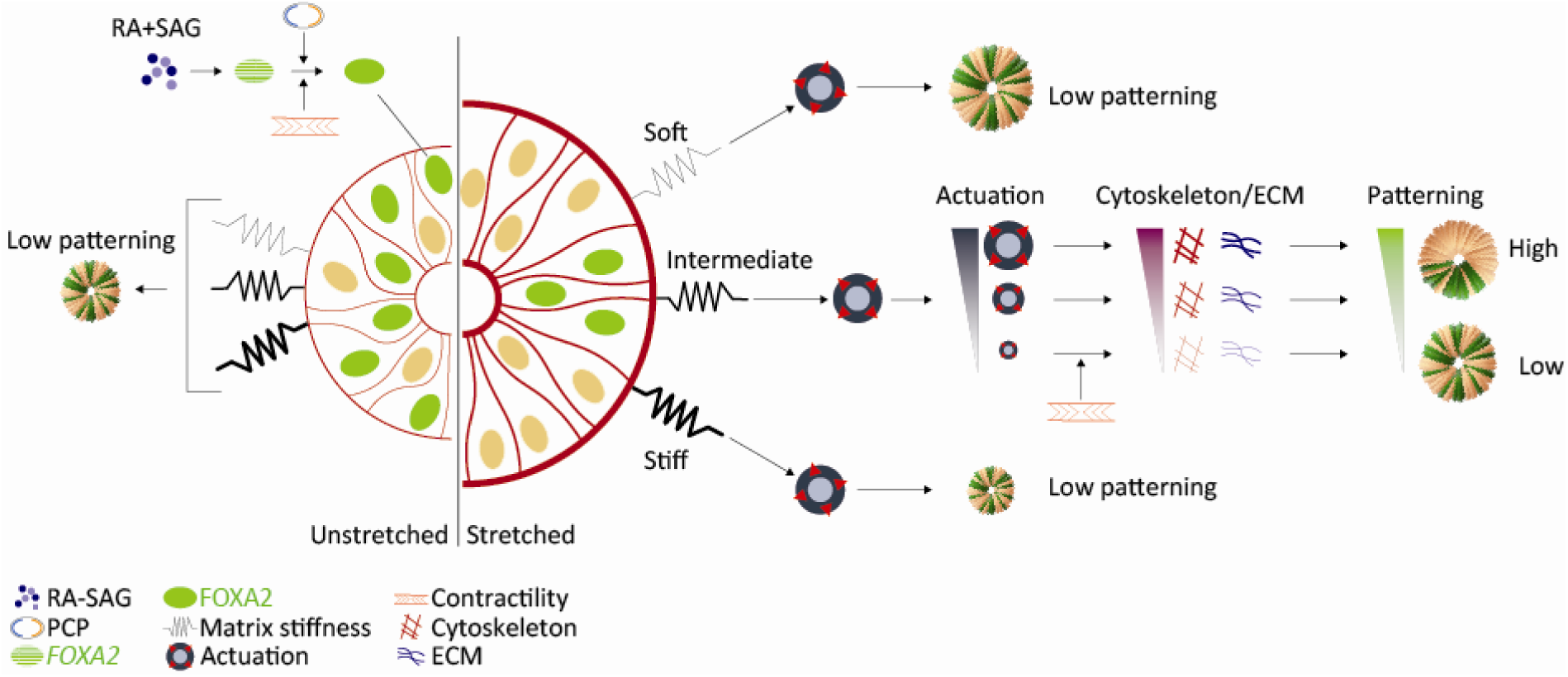
Proposed model for FP induction and patterning in hNTOs. Induction of the FP requires active contractility and PCP. FP patterning enhancement relies on hNTO actuation in a matrix stiffness-dependent manner. Soft and stiff matrices result in no FP patterning enhancement upon actuation. In intermediate stiffness, actuated hNTOs exhibit higher FP patterning events linked to increased cytoskeleton and ECM remodeling activity.

## Acknowledgements

This work was supported by the FWO grant G087018N and FWO postdoctoral fellowship 1217220N, Interreg Biomat-on-Chip grant and Vlaams-Brabant and Flemish Government co-financing, KU Leuven grants C14/17/111 and C32/17/027 and King Baudouin Foundation grant J1810950-207421. RF and YL were supported by NIH grants HD095520 and HD081216.

We thank Gert Hulselmans, Jasmien Willems and Vincent Merens for technical assistance.

## Author Contributions

AA conducted experiments and analysis. BD, GR, MB and SP conducted experiments. AA, BG and DR developed the mechanical stretching instruments and associated protocols. YL, XC and RF characterized the PCP-KO line. AA, SP, KD and SA generated and analyzed the scRNAseq data. AA, RF, MS, CV, HO, PD, SA and AR interpreted the data and edited the manuscript. AA and AR wrote the manuscript.

## Competing Interests

RF was CEO of Teratomic Consulting, LLC, which has been dissolved. He is also an associate editor of the journal Reproductive and Developmental Medicine, which provides his travel expenses.

## Materials and Methods

### Human PSC lines

The human PSC lines used in this study are 1) ZO1 hIPSC line (Mono-allelic mEGFP-Tagged TJP1 WTC hPSC Line, Coriell institute for Medical Research), 2) hIPSC line reported in Sahakyan, V. et al. Sci Rep 8, 018-21103 (2018), and 3) NCRM-1 (RRID:CVCL_1E71) hIPSC line from NIH Center for Regenerative Medicine (CRM), Bethesda, USA.

### Culture medium

Essential 8 (E8) - Flex Medium Kit (ThermoFisher Scientific) supplemented with 1% Penicillin Streptomycin (GIBCO) was used to maintain hPSC culture. The neural differentiation medium comprised of 1:1 mixture of neurobasal medium (GIBCO) and DMEM/F12 (GIBCO), 1% N2 (GIBCO), 2% B-27 (GIBCO), 1 mM sodium pyruvate MEM (GIBCO), 1 mM glutamax (GIBCO), 1 mM non-essential amino acids (GIBCO) and 2% Penicillin Streptomycin (GIBCO).

### Human PSC culture

Human PSCs were cultured in Matrigel-coated 6 well plates to 60-70% confluence. To passage the colonies, a 4 min treatment of Dispase (Sigma) was applied at 37°C, followed by three PBS washes. 1 mL of E8-Flex medium was added and the colonies were scraped and gently agitated using a pipettor to break the colonies. The colonies were then split 1:6 and incubated in 2 mL of E8Flex medium supplemented with Y-27632 Rock inhibitor ROCKi (Hellobio) at 10 μM for 24 h. The medium was then replaced by 4 mL of ROCKi-free E8-Flex medium and incubation was resumed for 48 h, at which point the colonies reached 60-70% confluence and were ready for passaging.

### Human NTO culture in PEG hydrogels

At a confluence of 60-70%, hPSCs were washed 3 times with PBS and dissociated into single cells by incubation in 1 mL of TrypLE Express (GIBCO) at 37°C for 3 min. Subsequently, 9 mL of DMEM/F12 media supplemented with 20% FBS (GIBCO) was added and cells were centrifuged at 300 RCF for 3 m. The supernatant was discarded and replaced by a second wash of 10 mL of DMEM/F12 media containing 20% FBS. Cell count was performed and remaining cells were centrifuged again at 300 RCF for 3 m. The pellet was resuspended in neural differentiation medium containing 10 μM of ROCKi to obtain a cell density of 3.5 M cells per mL. The cells were added to a nd-PEG premix to constitute 10% of the hydrogel’s total volume (see hydrogel preparation). 15 μL droplets of cell-hydrogel mix were placed on the surface of elastomeric membranes that underwent a silanization process through a nitric acid and (3-Aminopropyl)triethoxysilane (Sigma-Aldrich) treatment, and were mounted on equibiaxial or uniaxial stretching devices. Control membranes were placed inside a 10 cm culture dish. After gelation was visibly confirmed, an additional 20 m waiting time ensured complete gelation, and was followed by the addition of 2.5 mL of neural differentiation medium supplemented with 10 μM of ROCK inhibitor. After 72 hours the medium was replaced with 7 mL of neural differentiation medium supplemented with retinoic acid (RA) (Stemcell Technologies) at 0.25 nM and smoothened agonist (SAG) (Stemcell Technologies) at 1 μM. The medium was fully refreshed every 2 days until endpoint at day 11. Increasing the medium volume from 2 to 7 mL ensured that media levels did not drop below the height of the PEG droplets upon stretch.

To change the RA-SAG treatment window along the experimental timeline, the treatment was applied for days 1-3, days 3-5, days 5-7, days 7-9 and days 9-11 using a 96 well plate with 5 μL hydrogel droplets and 200 μL medium volume, following a 2 day media change protocol except for the earliest treatment. To apply prolonged ROCKi treatments, ROCKi was added to the medium at 10 μM at day 0 and supplemented to all media changes until day 11.

### PEG hydrogel preparation

PEG hydrogel precursors and buffers were prepared as previously described^24,25^. In brief, 8arm-PEG-vinylsulfone, 40 kDa (PEG-VS) (NOF Corporation), was functionalized with FXIIIa-peptide substrates via Michael type addition. We have used a glutamine-containing peptide (NQEQVSPLERCG-NH2) and lysine-containing peptides with a non-MMP-insensitive i.e. non-degradable sequence (AcFKGG-GDQGIAGF-ERCG-NH2). We obtained a glutamine-PEG precursor and a lysine-PEG precursor. Peptides were added to PEG-VS in a 1.2-fold molar excess over VS groups in 0.3 M triethanolamine (pH 8.0) at 37 °C for 2 h, followed by dialysis (Snake Skin; MWCO 10k; Pierce) against ultrapure water for 4 d at 4 °C. After dialysis, the salt-free products were lyophilized to obtain a white powder. These functionalized PEG powders were then reconstituted in water [10% (wt/vol)] to obtain working stock solutions of PEG.

Factor XIII (FXIII) was reconstituted in water and activated with thrombin: 1mL of FXIII (200 U/mL) and Fibrogammin P1250 (CSL Behring) were activated in the presence of 2.5 mM CaCl2 with 100 μL of thrombin (20 U/mL; Sigma-Aldrich) for 30 min at 37 °C. Aliquots of FXIIIa were stored at –80 °C for further use. Precursor solutions to give hydrogels with a final dry mass content as indicated in the table below were prepared by stoichiometrically balanced ([Lys]/[Gln] = 1) solutions of glutamine-PEG and lysine-PEG in Tris buffer (TBS; 50 mM; pH 7.6) containing 50 mM calcium chloride. The cross-linking reaction was initiated by 10 U/mL thrombin-activated FXIII and vigorous mixing.

The hydrogel composition is described in the table below and provides matrices with varying stiffnesses.

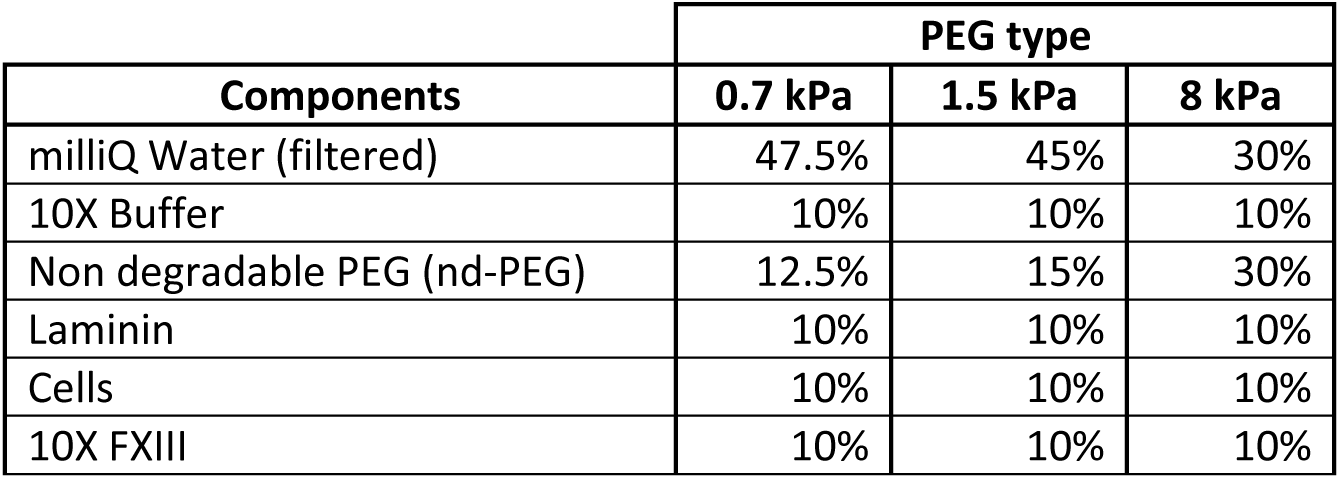

| Components | PEG type |  |  |
| --- | --- | --- | --- |
|  | 0.7 kPa | 1.5 kPa | 8 kPa |
| milliQ Water (filtered) | 47.5% | 45% | 30% |
| 10X Buffer | 10% | 10% | 10% |
| Non degradable PEG (nd-PEG) | 12.5% | 15% | 30% |
| Laminin | 10% | 10% | 10% |
| Cells | 10% | 10% | 10% |
| 10X FXIII | 10% | 10% | 10% |

First, milliQ water, 10X Buffer and nd-PEG were mixed and kept at room temperature, until the hPSCs were prepared for embedding. After the addition of laminin (BD) to a final concentration of 0.1mg/ml and 10X FXIII, the premix was ready for immediate cell embedding followed by plating. We have previously shown that laminin is covalently incorporated into the FXIII-crosslinked PEG and that the addition of 0.1mg/ml laminin does not alter the mechanical properties of this matrix^24^.

To supplement the hydrogels with red fluorescent beads (FluoSpheres Carboxylate-Modified Microspheres, 200 nm, Invitrogen), the stock solution is first diluted 1:10 before adding it to the hydrogel premix for a final composition of 2% (1:500 dilution of the stock solution). The volume of fluorescent beads added is subtracted from the volume of milliQ water in the premix to ensure the total composition of the hydrogel is not compromised.

### Stretching Devices

An equibiaxial stretching device was used to apply equibiaxial strains on compatible elastomeric membranes hosting the hNTO containing hydrogels. Stretch was applied through the rotation of 8 arms that connect to the membranes. The motor unit and the setup were placed in the incubator. The stretching schedule was controlled through an external computer. This allowed the modulation of stretch magnitude and duration.

A uniaxial stretching was also used to provide linear strains on the same membranes as the equibiaxial device. The apparatus connected to the membrane at two ends, one fixed and one free-moving. A motor unit translated the free-moving end applying linear stretching. The entire instrument was placed in the incubator except for the programmable logic controller (ArduinoUno) which controls the rate and timing of stretching and remains outside the incubator.

### Immunohistochemistry

4% paraformaldehyde (Sigma-Aldrich) was used to fix hNTOs for 2 hours followed by washing with PBS. Permeabilization and blocking solutions were made with 0.3% Triton X (PanREAC AppliChem) and 0.5% BSA solution (Sigma-Aldrich) for 30min. Primary antibodies (**Table S3**) were suspended in permeabilization and blocking solution and applied to the fixed and permeabilized hNTOs for 24 h. Three PBS washes were performed over an additional 24 h period. Subsequently, immunolabeling was performed over a duration of 24 h using donkey-anti mouse Alexa Fluor 555/647 (Invitrogen), donkey-anti rabbit Alexa Fluor 555/647 (Invitrogen), donkey-anti goat Alexa Fluor 555/647 (Invitrogen) and donkey-anti rat Alexa Fluor 647 (Jackson ImmunoResearch) secondary antibodies. To visualize filamentous actin we employed Alexa Fluor 647 conjugated phalloidin (Abcam). Hoechst was used to visualize nuclei. Click-it EDU Alexa Fluor 647 kit (Invitrogen) was used to label cells with active DNA synthesis. To visualize ZO1, the ZO1 reporter cell line was used.

### Image acquisition and analysis

For quantification analysis, fluorescence images were obtained using an inverted microscope (Zeiss Axio Observer Z1; Carl Zeiss MicroImaging) equipped with a Colibri LED light source and a 10x air objective. Confocal representative images were obtained using a confocal microscope (Leica SP8 DIVE, Leica Microsystems) operated on either confocal or multiphoton mode with 25x water objective. Live imaging was performed using an inverted microscope (EVOS, Invitrogen) using a 4x and 10x air objectives under transmitted light or using an RFP filter for bead tracking experiments.

Possible differences in the stress states within the hydrogel along vertical axis are small as demonstrated by simulation (**Supplementary Fig. 5**). However, to avoid any potential variations in stress, organoid settling within the hydrogel prior to polymerization restricted hNTO vertical positions to the bottom-most region, and therefore closest to membrane surface. This ensured that all analyzed hNTOs were subjected to comparable stress states when stretched and that the imposed strains most reflected that of the membrane.

### Organoid size evaluation

To evaluate hNTO size, bright field images were analyzed using ImageJ (Version 1.52a) by manual tracing of organoid borders to obtain an area which is then converted to an equivalent diameter.

### Patterning assessment in hNTOs

To evaluate FP domain size, fluorescence images were analyzed using ImageJ by manually tracing regions that span FOXA2+ cells within hNTOs resulting in a FOXA2+ area. To obtain FOXA2 AR, FP domain size was divided by hNTO area. To systematically determine the scattering or patterning of FP domain within an hNTO, 1) the FOXA2 AR was determined, and 2) a FOXA2 intensity ratio (FIR) was obtained where the average FOXA2 intensity of the FOXA2+ region was first subtracted by the average FOXA2 intensity over the entire hNTO area, then divided by the latter. Subsequently, an hNTO was evaluate as scattered for 1) AR > 0.5, and 2) AR < 0.5 but FIR < 20%. By contrast, an hNTO was evaluated as patterned for AR < 0.5 and FIR > 20%. These criteria were determined upon visual investigation of multiple FOXA2 expressions. To evaluate positional information of labeled samples, the line section in ImageJ was used. Although area quantification is used here, by using wide field microscopy we capture the majority of signal from the organoid volume allowing us to distinguish between patterned and scattered expressions of FOXA2.

### Bead tracking experiments

Bead tracking experiment involved tracking fluorescent beads embedded within the matrix. Imaging was performed every 2 days starting from day 1. To analyze matrix strain and stress states, ImageJ was used where, at day 1, a segment normal to the border of the hNTO at its widest projected area, in the radial direction of stretching, and linking the hNTO border to a bead in the same focal plane is first identified, signifying a reference length. For every subsequent day, the reference length is subtracted from the newly measured segment length and the result is divided by the former to evaluate the strain component in the direction of stretching ε_rr_. The matrix along the stretch direction is considered 1) in tension for ε_rr_ > 0, 2) in compression for ε_rr_ < 0, and 3) neutral for ε_rr_ = 0. The ε_rr_ values were converted to the stress component in the direction of stretching σ_rr_ = Eε_rr_ using matrix stiffness values E. Although this is a simplified stress interpretation from a 2D evaluation of a 3D environment, it nevertheless underscored a general stress-growth rate relationship. For each hNTO, 5 equally spaced segments around the organoid are observed and analyzed to provide an average stress state for the surrounding matrix. To determine the critical stress point σ_rr,critical_ (**Fig. 3b**), the MATLAB (R2018a, The MathWorks Inc.) Knee Point function was used.

### PEG hydrogel equibiaxial simulations

To simulate equibiaxial stretching of 2 kPa PEG hydrogels Abaqus (Dassault Systemes, 2019) was used. To represent the PEG matrix droplet, a 100 units by 25 units curved part was created in 2D axisymmetric configuration with elastic modulus E = 2,000 Pa and Poisson’s ratio of 0.5. A 100 units by 10 units part representing the membrane was linked and fixed to the bottom surface of the matrix with a similar E. One side of the membrane was displaced by 22 and 41 units representing a strain ε_rr_ = 0.22 and 0.41 as a results of increasing the membrane area by 50% and 100%. The membrane part was hidden for visualization purposes (**Supplementary Fig. 5**).

### Human NTO isolation and single-cell RNA sequencing

Organoids were grown following the hNTO differentiation protocol under five conditions 1) equibiaxially stretched – Day 11, 2) unstretch– Day 11, 3) equibiaxially stretched – Day 5, 4) unstretched – Day 5, and 5) hNTOs grown for 3 days (before stretching and RA/SAG treatment). For dissociation, the PEG gels were gently detached from the elastic membranes with a pipette tip and transferred to a clean and dry petri dish. A scalpel was used to slice the 15 uL gel droplet into four to six pieces. A total of five 15 μL gel droplets were collected, processed and pooled as described for each condition. Pooling was done to ensure enough cells were present for the library preparation and sequencing steps. Gel fragments were transferred to a 15 mL falcon tube containing 1 mL of prewarmed TrypLE Express at 37°C. The samples were transferred to a warm water bath at 37°C for 7.5 minutes with gentle agitation every 1 minute after the first 3 minutes. Gradually the gel fragments, as well as visible organoids, began to visually disappear. A visual inspection was performed using an inverted microscope to ensure the presence of single cells. The 1 mL solution was introduced to 9 mL of DMEM supplemented with 20% FBS for TrypLE Express neutralization, and centrifuged at 500 RCF for 5 minutes. The pellet was resuspended in 200 uL of N2B27 media and put on ice. This optimized dissociation protocol yielded an average viability of 85.4% across both stretched and control samples.

Library preparations for the scRNAseq was performed using 10X Genomics Chromium Single Cell 3’ Kit, v3 (10X Genomics, Pleasanton, CA, USA). The cell count and the viability of the samples were accessed using LUNA dual fluorescence cell counter (Logos Biosystems) and a targeted cell recovery of 6000 cells was aimed for all the samples. Post cell count and QC, the samples were immediately loaded onto the Chromium Controller. Single cell RNAseq libraries were prepared using manufacturers recommendations (Single cell 3’ reagent kits v3 user guide; CG00052 Rev B), and at the different check points, the library quality was accessed using Qubit (ThermoFisher) and Bioanalyzer (Agilent). With a sequencing coverage targeted for 50,000 reads per cell, single cell libraries were sequenced either on Illumina’s NovaSeq 6000 or NovaSeq 500 platforms using paired-end sequencing workflow and with recommended 10X; v3 read parameters (28-8-0-91 cycles). The data generated from sequencing was demultiplexed and mapped against human genome reference using CellRanger v3.0.2.

### Single Cell RNA sequencing Data Processing

We have sequenced 4,694, 2,541, 4,687, 6,320 and 5,708 cells for day 3, unstretched day 5, stretched day 5, unstretched day 11 and stretched day 11 samples respectively for a total of 23,950 cells which were reduced after QC steps to 17,826 cells with an average of 4,147 224 detected genes per cell. Data manipulation and subsequent steps were performed using the Seurat^51^ tool for single cell genomics version 3 in R version 3.4. A filtering step was performed to ensure the quality of the data, where the counts of mitochondrial reads and total genes reads was assessed. Cells with more than 15% of identifiable genes rising from the mitochondrial genome were filtered out. Similarly, cells having fewer identifiable genes than 200 (low quality) and above 7,500 (probable doublets) were filtered out. Data normalization was performed, followed by the identification of 2,000 highly variable genes using the FindVariableFeatures. S-phase and G2M-phase cell cycle regression was performed to allow cell clustering purely on cell identity and fate, which otherwise was biased by cell cycle phases. Auto scaling of the data was performed and described using principal component analysis (PCA) using the RunPCA function.

### Data Clustering

Graph-based clustering using the FindNeighbors function (using top 15 principal components (PCs)) and FindCluster function (resolution = 0.5) was performed to group cells based on their transcriptional profiles. No batch correction was performed on the data set. Data visualization was performed using the Uniform Manifold Approximation and Projection (UMAP) dimensionality reduction technique using the RunUMAP package while employing the top 15 PCs identified in the previous PCA step. Cluster annotation was aimed at identifying D-V regions based on the expression level of several hallmark genes related to anteroposterior (A-P) as well as dorsoventral (D-V) regionalization. The hallmark genes were grouped to create gene-sets for Forebrain (FB) (*FOXG1, LHX2, DLX2, NKX2*.*1, GSX2, SIX3*), Midbrain (MB) (*PAX5, LMX1A, LMX1B, SIM1, EN1, EN2*), Hindbrain (HB) (*GBX2, HOXA2, HOX4, HOXB2, HOXB4*), Dorsal (D) (*LMX1A, OLIG3, PAX3, PAX7*), Intermediate (I) (*PAX6, DBX1, DBX2*), and Ventral (V) (*NKX6*.*1, OLIG2, NKX2*.*2, FOXA2*). Using these gene-sets, we performed the following cluster identification: D-11 (dorsal with hindbrain, day 11), I-11 (intermediate with midbrain, day 11), FB (forebrain and mostly ventral, day 11), V-11 (ventral and mixed A-P, day 11), V-5 (ventral with forebrain and midbrain, day 5), NP-5 (neuroprogenitors, with forebrain and midbrain, day 5), and NP-3 (neuroprogenitors with forebrain and midbrain, day 3). These gene-sets were used to obtain a score for every cell using the AddModuleScore function. Other markers were included to identify additional clusters that did not exhibit the abovementioned gene-sets, such as Neural Crest (NC) markers (*SOX 10, MPZ, FOXD3*), Neural Crest Derivative cells (NCD) (*TWIST1, TWIST2, FOXC1*). These markers were used to identify the NC (mixed days) cluster as well as broadly define the NCD (day 11) cluster which is a merger of 3 sperate subclusters. Clusters that did not have particular strong expression of the above markers were designated transition clusters (T-a, and T-b with mixed days) as they seemed to link the data sets of days 3 and 5 to those of day 11. As such, these clusters were enriched in multiple markers, for example neuronal markers *ISL2* as well as important dorsal signaling marker *BMP4* in T-a and T-b, *FOXA2* in T-b, and *HOXB4* in T-a. One cluster was manually removed due to very low unique molecular identifier (UMI) count, after which the data was reprocessed to account for the removal of the cluster.

### SCENIC

Transcription factor network inference was performed using the SCENIC^40^ pipeline on the combined scRNAseq dataset. Regulon network activity was evaluated and represented by the scoring step AUCell for each cell using default settings on the full dataset. The AUC values for the FOXA2 regulon were implemented in the Seurat data matrix file and used for visualization using UMAPs.

### Pseudotime trajectories

Trajectory analysis was performed using Monocle 2^52^. We performed the analysis on the BIOMEX platform (VIB-KU Leuven Center for Cancer Biology, BIOMEX) by importing the Seurat obtained normalized count matrix, metadata and using the 2,000 Seurat obtained highly variable genes under default settings.

### Correlation analysis

Correlation heatmap between stretch conditions was performed on the BIOMEX platform using the Seurat obtained normalized count matrix, metadata and using the 2,000 Seurat obtained highly variable genes under default settings.

### D-V cell binnning

To explore cell abundance along the D-V axis, we assigned each cell an A-P and a D-V position using the combined data set. Through this cell binning step, the A-P assignment was based the FB, MB, and HB gene-sets, while the D-V one was based on the D, I and V gene-sets. Using the AddModuleScore function, a cell was assigned the A-P position (FB, MB or HB) that scored highest for the respective gene-set. In a similar manner the D-V position (D, I and V) was assigned. We only retained cells with positive assigned A-P and D-V gene-set scores and only those that could be assigned a position on both axes. Accounting for the A-P score ensured the D-V assignment was biologically relevant. Finally, to obtain the D-V fractions we divided the amount of cells belonging to each assigned position (D, I or V) by the total cells along the D-V axis. The process was conducted for samples of days 5 and 11 for both stretch conditions.

### Differential gene expression (DGEA) and gene ontology (GO) enrichment analysis

To perform a DGEA we employed the FindMarkers function, a part of the Seurat R package, between all day 11 cells, with function parameters min.pct = 0.0 and logfc.threshold = 0.0 to capture the complete gene list across the full range of LFC.

To perform the GO enrichment analysis, we first extracted the full list of differentially expressed genes ranked by LFC from highest to lowest. This list was then inserted into the GOrilla^53^ web portal using the Homo sapiens organism, single ranked file running mode, and a p-value threshold of 10^−4^. The results were then passed to the REVIGO^54^ web portal to obtain a summarized list of gene ontology terms using the default setting and small similarity option. The gene ontology terms were then ranked by significance of which relevant processes were reported.

### Quantification and Statistical Analysis

For statistical analysis we employed a two-way ANOVA statistical test on grouped data, and an unpaired two-tailed t-test where appropriate with a 95% confidence interval (GraphPad Prism 6, Version 6.01, GraphPad Software, Inc.). When determining patterning and scattering significance between conditions, the statistical analysis was performed using the patterned values of the various conditions. Similarly when determining the FOXA2+/-hNTOs, the statistical analysis was performed using the FOXA2+ hNTO values of the various conditions. Pearson correlations were performed to evaluate linear regression where appropriate. Statistical significance was considered for all comparisons with p < 0.05. For hierarchical clustering, we employed the R package heatmap.2.

### Data availability

All raw sequencing data of the scRNAseq experiments, and the combined processed and metadata files generated in this study will be made available at GEO.

