## Supplementary Information for "Actuation Enhances Patterning in Human Neural Tube Organoids"

**Supplementary Figures**

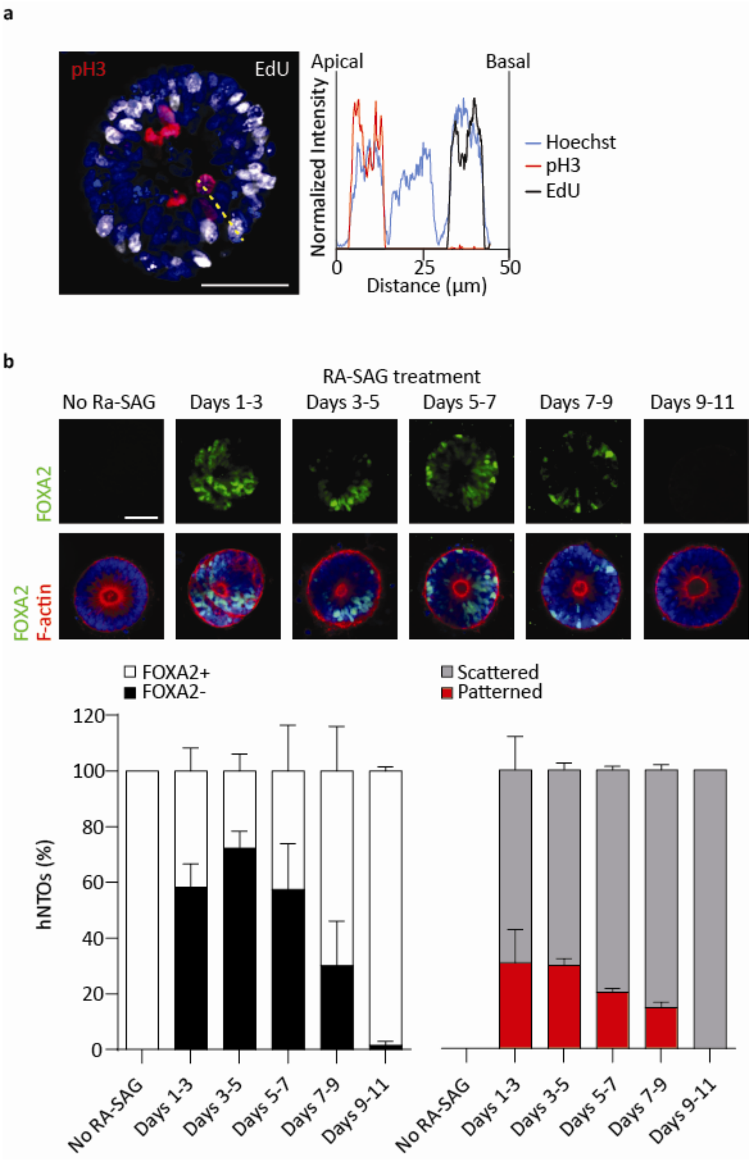


**Supplementary Fig. 1 a** Representative hNTO at day 11 hNTO displaying mitotic cells (pH3) near the apical side and S-phase cells (EdU) near the basal side. A 50 µm line section shows the normalized intensity profile of pH3 (mitotic cells, red), EdU (S-phase, black), and Hoechst (nuclei, blue), suggesting interkinetic nuclear migration characteristic of pseudostratified epithelia. **b** FP induction and patterning frequencies upon RA-SAG treatments shifted along the experimental timeline in 2-day increments. FP expression is evaluated at day 11 on fixed and permeabilized hNTOs (n = 3 except for RA-SAG treatment days 1-3 (n = 2) for > 50 hNTOs per condition). Error bars are SEM, scalebars 50 µm.

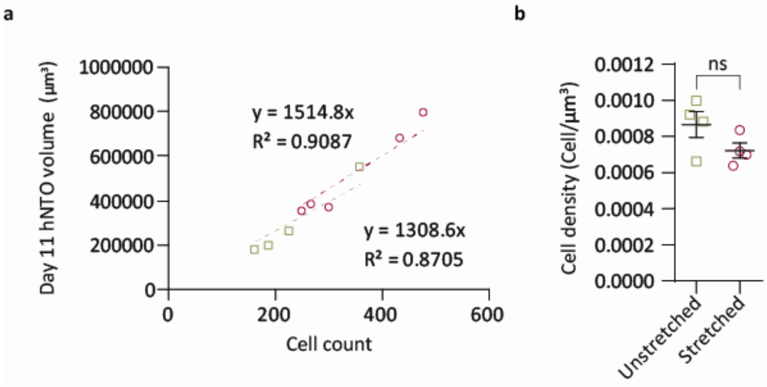


**Supplementary Fig. 2** Cell count in unstretched and stretched hNTOs. **a** Relationship between total cell count and organoid volume (n = 4 for unstretched hNTOs and n = 5 for stretched hNTOs). **b** Calculated cell density in unstretched and stretched organoids (n = 4 unstretched hNTOs and stretched hNTOs).

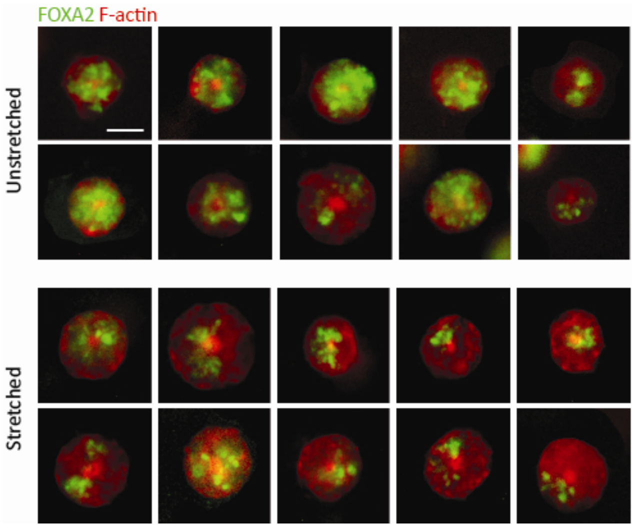


**Supplementary Fig. 3** Representative images of hNTOs stained for FOXA2 and F-actin in unstretched and stretched conditions. FOXA2 is more abundant and scattered in unstretched hNTOs as measure by $AR_{avg}$ = 0.46 compared to stretched ones with AR = 0.29 that display a higher degree of patterning events. Scalebar 50 µm.

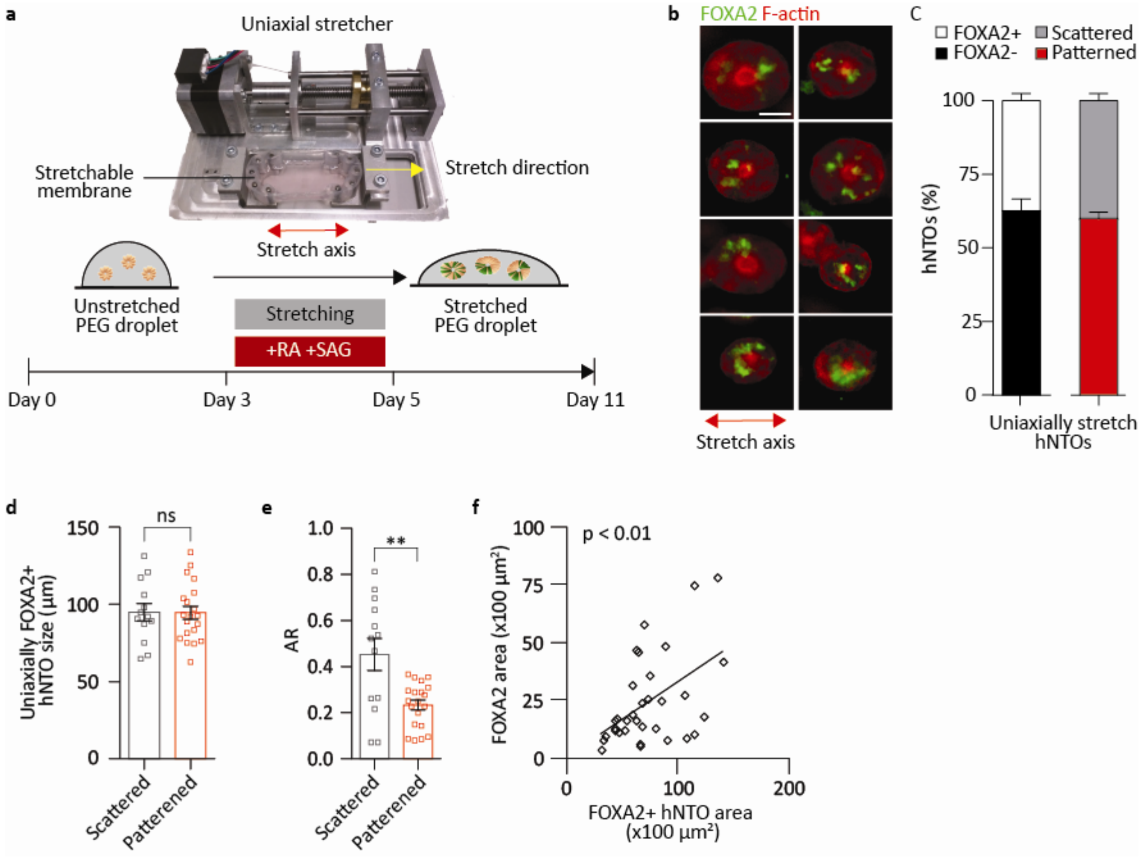


**Supplementary Fig. 4** Uniaxial stretching of hNTOs increases FP patterning events. **a** PEG hydrogel droplets are placed onto stretchable membranes attached to a uniaxial stretching device. Stretching occurs simultaneously with the RA-SAG treatment (days 3-5) and membrane length increases along the stretch axes by 20%. The strains are transferred to the PEG hydrogel and in turn transmitted to the embedded hNTOs. **b** Representative images displaying FOXA2 and F-actin expressions in stretched conditions. hNTOs are elongated along the stretch axis as indicated by the red arrow. **c** Quantification of FP induction and patterning for uniaxially stretched samples. (n = 3 for a total of 50 hNTOs). **d** Comparison of patterned and scattered organoid size in uniaxially strained hNTOs (Pooled FOXA2+ hNTOs from **c**). **e** Quantification of AR for scattered and patterned hNTOs upon uniaxial stretch (pooled FOXA2+ hNTOs from **c**). **f** Scatter plot of FOXA2 domain area for corresponding organoid area shows FP domain scaling with hNTO size (Pooled FOXA2+ hNTOs from **c**). Uniaxial straining imparts similar FP behavior as equibiaxially stretched hNTOs. Error bars are SEM, ** p<0.01, scalebar 50 µm.

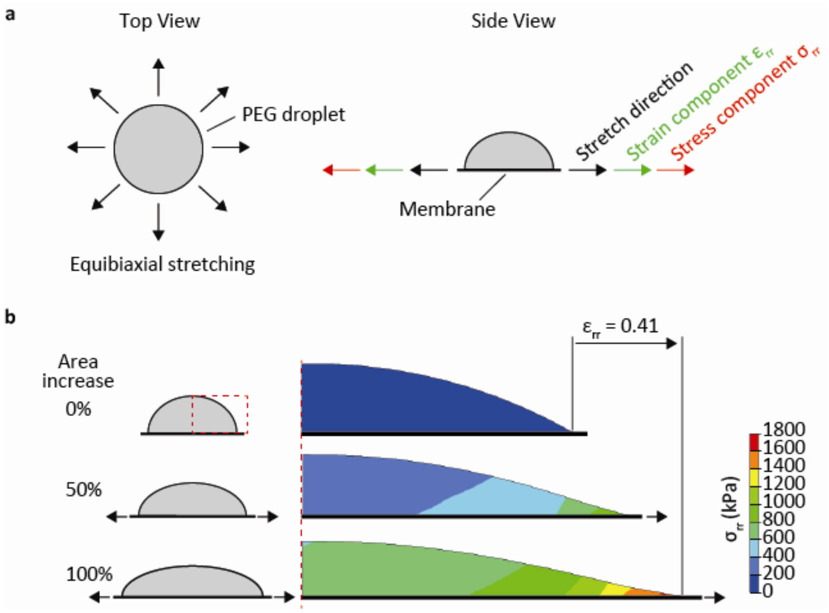


**Supplementary Fig. 5 a** Top view of equibiaxial stretching of a PEG droplet. Black arrows showing stretch direction. Side view of the same droplet showing stretch direction and associated strain component $\varepsilon_{rr}$ (green arrows) and stress component $\sigma_{rr}$ (red arrow) in the direction of stretching. **b** Axisymmetric equibiaxial stretching simulation of 2 kPa PEG hydrogel. Stress component values, $\sigma_{rr}$, are largely homogeneous throughout the PEG droplet except for the outermost regions. The inner regions have an average stress value of 760.2 ± 129.4 kPa, comparable to *in vitro* measured stretch-induced stresses of 637.3 ± 124.2 kPa. The simulation suggests that hNTOs in the bulk of the matrix will be subjected to comparable stress states.

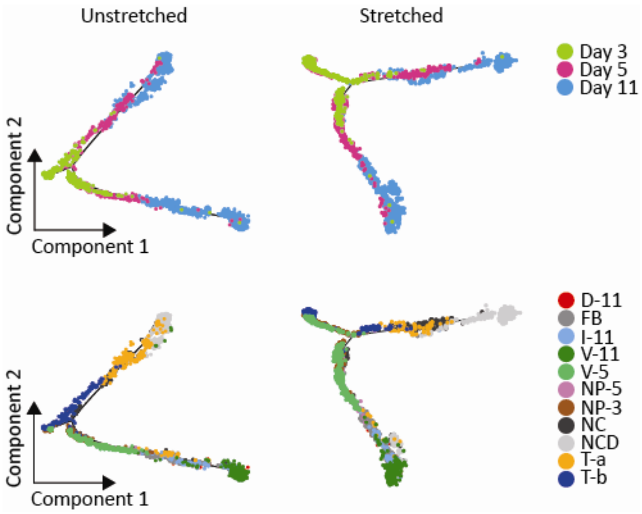


**Supplementary Fig. 6** Pseudotime trajectories for unstretched and stretched hNTOs color coded for days, and annotated clusters. Trajectories shows a progression from day 3 to day 11 with a bifurcation that leads to 1) neural fates and 2) neural crest and neural crest derivatives fates. Stretched hNTO pseudotime trajectory is comparable to the unstretched case.

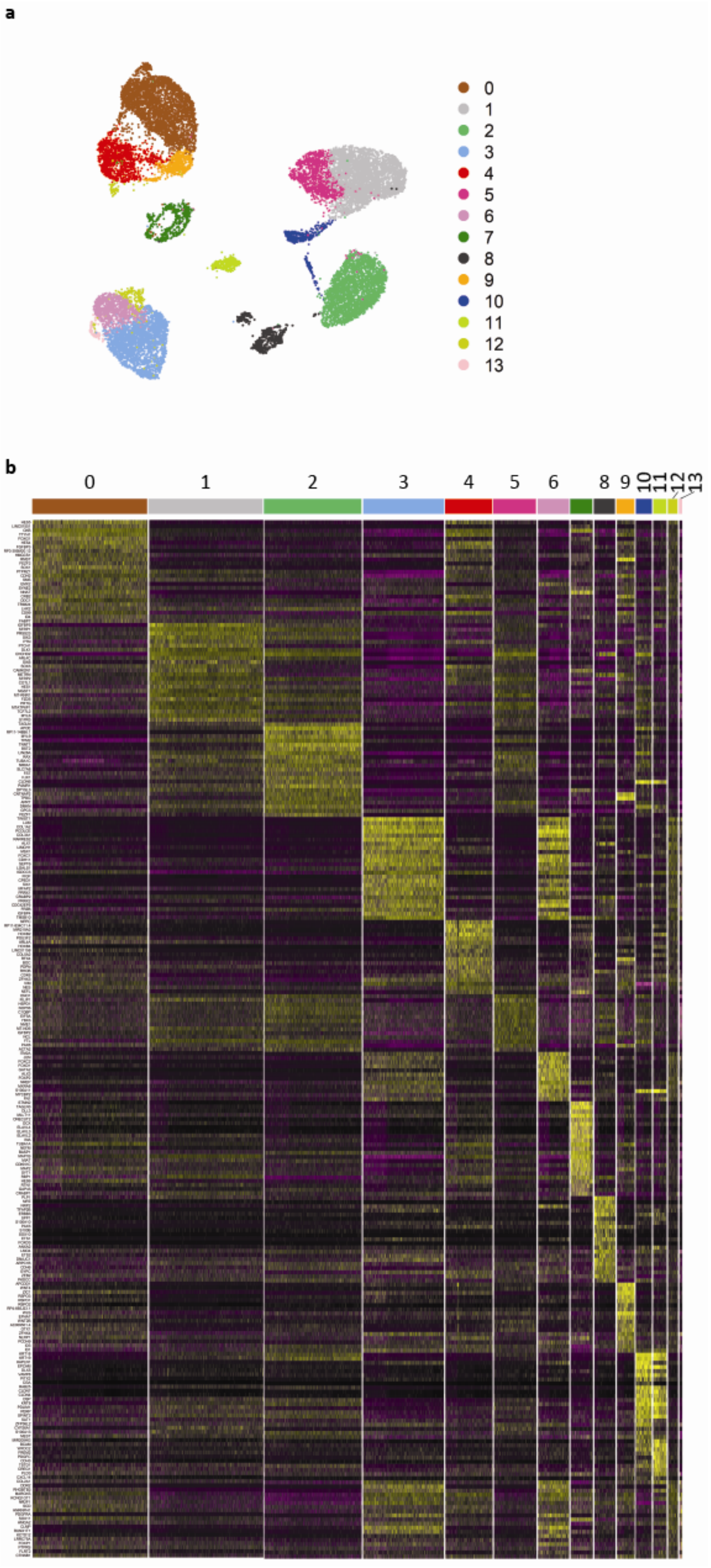


**Supplementary Fig. 7 a** Unannotated UMAPs showing 14 clusters for the combined day 3, 5 and 11 unstretched and stretched dataset. **b** Gene expression heatmap showing top 25 marker genes for each cluster.

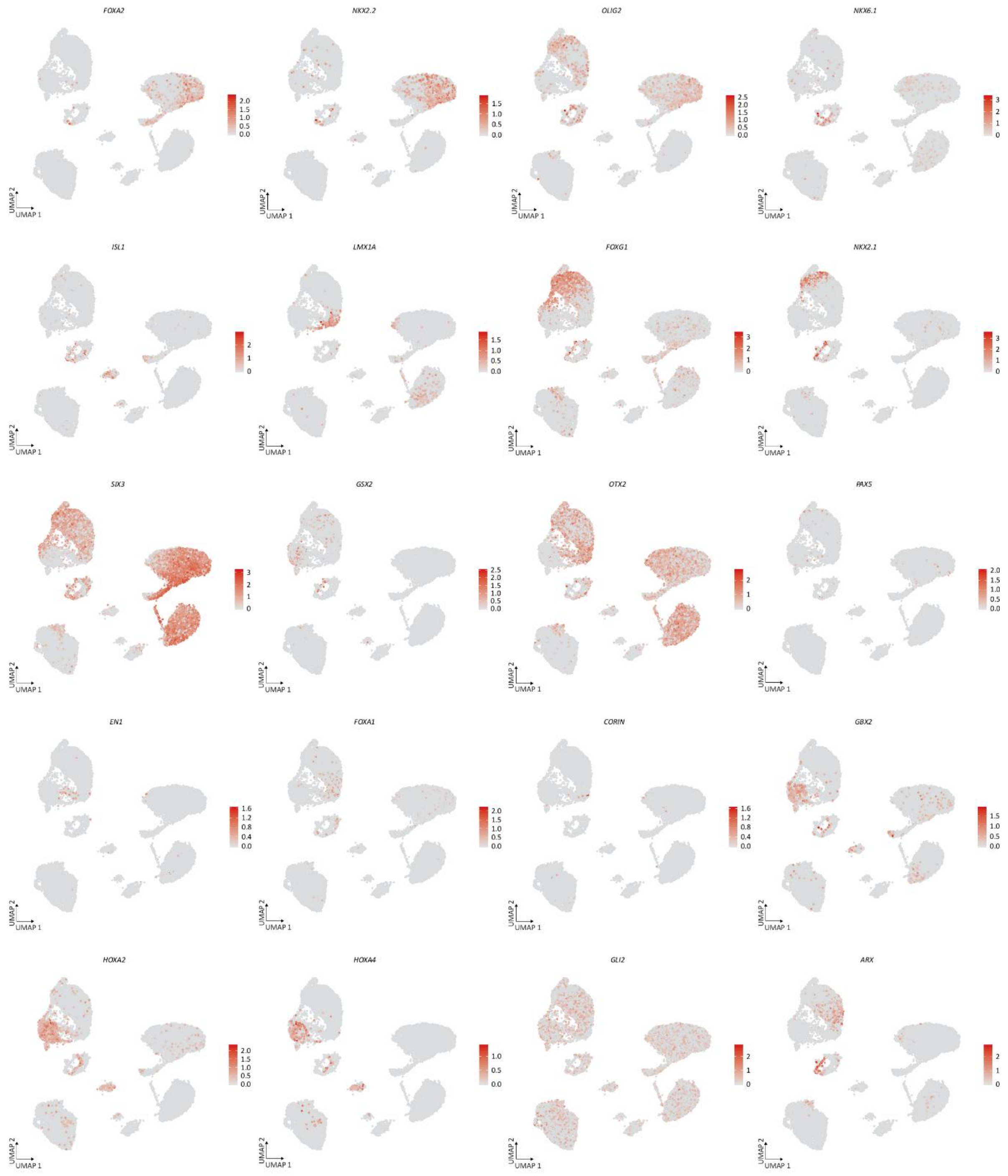


**Supplementary Fig. 8** Neural differentiation transcriptomic markers.

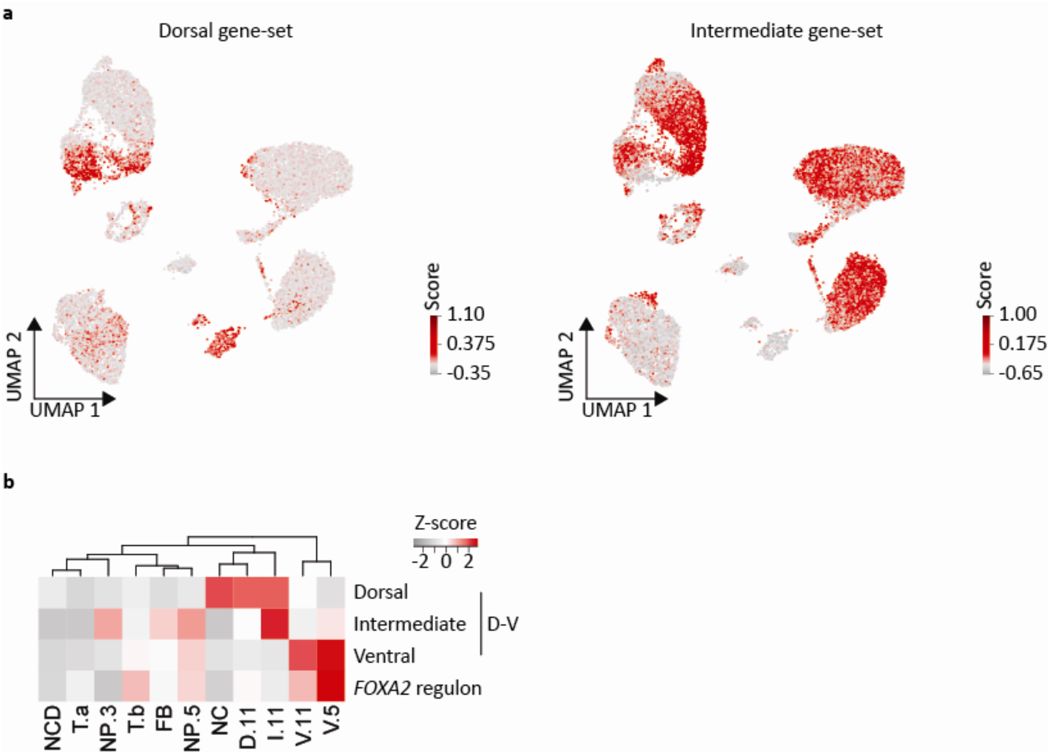


**Supplementary Fig. 9 a** UMAPs showing D-V (D and I, for V see **Fig. 4d**) gene-set scores. **b** Heatmap displaying hierarchically clustered clusters based on D-V gene-set scores and *FOXA2* regulon scaled AUC scores indicating that *FOXA2* regulon activity correlates with the ventral identity.

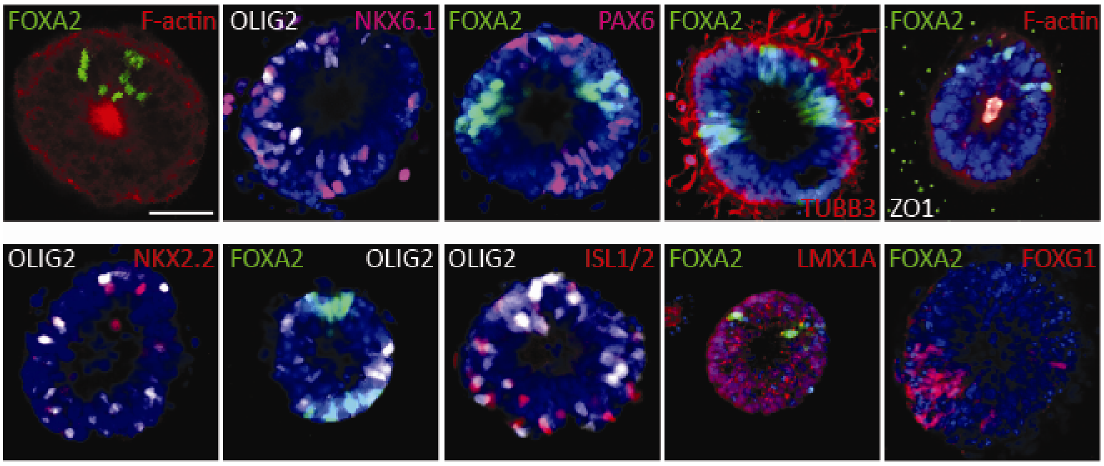


**Supplementary Fig. 10** Immunohistochemistry showing cell identities in stretched hNTOs. Scalebar 50 µm.

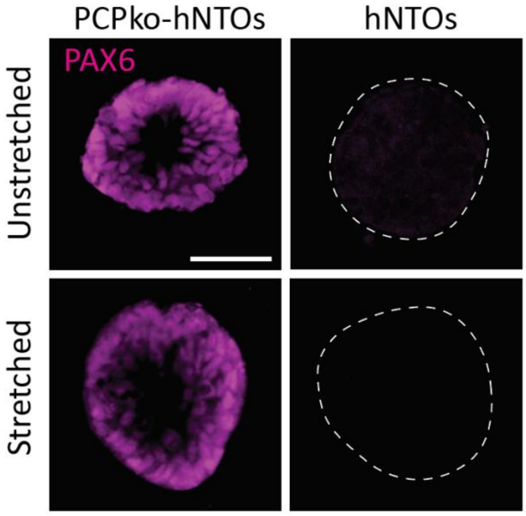


**Supplementary Fig. 11** Representative images displaying more abundant PAX6 expression in PCP$^{KO}$-hNTOs compared to control hNTOs under unstretched and stretched conditions. Scalebar 50 µm.

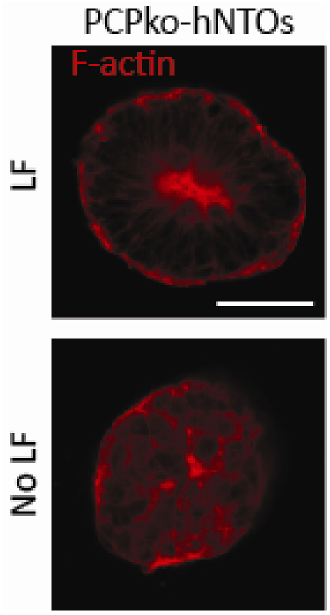


**Supplementary Fig. 12** Disrupted cytoskeleton organization in PCP$^{KO}$-hNTOs resulting in elongated lumens and organoids without LF. Scalebar 50 µm.

**Supplementary tables**

**Supplementary table 1.** Top 25 marker genes per annotated cluster

**Supplementary table 2.** Regulon AUC score per annotated cluster

**Supplementary table 3.** List of primary antibodies
